# Glutathione regulates transcriptional activation of iron transporters via *S*-nitrosylation of bHLH factors to modulate subcellular iron homeostasis

**DOI:** 10.1101/2021.05.09.443283

**Authors:** Ranjana Shee, Soumi Ghosh, Pinki Khan, Salman Sahid, Chandan Roy, Dibyendu Shee, Soumitra Paul, Riddhi Datta

## Abstract

While glutathione (GSH) is known to regulate iron (Fe) deficiency response in plants, its involvement in modulating subcellular Fe homeostasis remained elusive. In this study, we report that the GSH depleted mutants, *cad2-1* and *pad2-1* displayed increased sensitivity to Fe deficiency with significant down-regulation of the vacuolar Fe exporters, *AtNRAMP*3 and *AtNRAMP*4 and the chloroplast Fe importer, *AtPIC*1. Moreover, the *pad2-1* mutant accumulated higher Fe content in vacuoles and lower in chloroplasts compared with Col-0 under Fe limited condition. Exogenous GSH treatment could enhance the chloroplast Fe content in Col-0 plants but failed to do so in the *nramp3nramp4* double mutant demonstrating the role of GSH in modulating subcellular Fe homeostasis. Pharmacological experiments, mutant analysis and promoter assay revealed that this regulation involved transcriptional activation of the transporter genes by a GSH-GSNO module. The Fe responsive bHLH transcription factors (TFs), *At*bHLH29, *At*bHLH38 and *At*bHLH101 were identified to interact with the promoters of these genes which were in turn activated via *S*-nitrosylation. Together, the present study delineates the role of GSH-GSNO module in regulating subcellular Fe homeostasis by transcriptional activation of the Fe transporters *AtNRAMP3*, *AtNRAMP4* and *A*t*PIC1* via S-nitrosylation of the bHLH TFs during Fe deficiency.

**Summary statement:** Glutathione regulates subcellular iron homeostasis under iron deficiency via GSNO dependent transcriptional activation of *AtNRAMP3*, *AtNRAMP4* and *AtPIC1* genes by *S*-nitrosylation of the iron responsive bHLH transcription factors, *At*bHLH29, *At*bHLH38 and *At*bHLH101.

## Introduction

Iron (Fe) is an essential micronutrient for plant growth and development. It plays an important role in regulating numerous cellular responses because of its physicochemical properties. This micronutrient is known to coordinate metalloprotein active sites and regulate many important enzymatic reactions required for nitrogen fixation, DNA synthesis and biosynthesis of various phytohormones (Briat et al., 2015). Although Fe is found abundantly in the earth’s crust, it is usually present in an oxidized form and its availability to plants is limited. The deficiency of Fe in plants causes chlorosis and perturbs photosynthesis and oxidative phosphorylation by hampering different electron carriers of photosynthetic machineries as well as mitochondrial electron transport system. Therefore, Fe deficiency is a challenging issue for plant growth, development and productivity.

Plants have evolved highly sophisticated mechanisms for Fe uptake and transport. Two basic strategies for Fe uptake are described in the graminecious and non-graminecious plants (Morrissey and Guerinot, 2009). Among them, the strategy I or reduction based strategy is found in the non-graminecious plants like *Arabidopsis* where the plasmamembrane H^+^-ATPases (AHAs) release protons to increase rhizosphere acidification. This promotes the reduction of Fe^3+^ to the more soluble Fe^2+^ by ferric reductase oxidase 2 (FRO2) (Robinson et al., 1999). Iron regulated transporter 1 (IRT1), a member of the ZIP metal transporter family, imports the reduced Fe^2+^ into the root cells (Vert et al., 2002). Also, shoot specific FRO7 plays an important role in Fe delivery to chloroplasts. Besides, FRO3 and FRO8, present in the mitochondrial membrane, also regulate Fe homeostasis in mitochondria (Jeong and Connolly, 2009).

Modulating intracellular Fe homeostasis is crucial particularly under altered Fe conditions and involves several Fe transporters. A high-affinity Fe transporter, natural resistance against macrophage protein (NRAMP), having significant similarity with its mammalian counterpart, was identified in *A. thaliana* (Curie et al., 2000). Although their functional role in plants was not entirely revealed, complementation assays in yeast showed that these proteins were induced under Fe deficient condition. NRAMP 1, 3 and 4 represent multi-specific metal transporters localized in roots and leaves (Curie et al., 2000; Thomine et al., 2003; Lanquar et al., 2005). In Arabidopsis, NRAMP3 and NRAMP4 were identified to retrieve and export Fe from vacuoles to chloroplast during the Fe deficient condition (Bastow et al., 2018). In fact, the vacuolar Fe released by these two transporters was found to be the primary source of Fe in germinating seeds (Bastow et al., 2018). On the other hand, vacuolar iron transporter 1 (VIT1) was identified as one of the key transporters involved in Fe influx into the vacuole. It thus functions in the vacuolar sequestration process essential for detoxification under the excess Fe condition (Kim et al., 2006). In plants, maintaining the plastidal Fe homeostasis is crucial for survival. Several Fe transporters like permease in chloroplasts 1 (PIC1), non-intrinsic ABC protein 11 (NAP11) and NAP14 were identified as Fe importers that transport Fe across the chloroplast envelope (Duy et al., 2007; Shimoni-Shor et al., 2010). On the other hand, several transporters like yellow stripe like 1 (YSL1) and YSL3 help in maintaining Fe homeostasis by regulating Fe efflux from the chloroplast (Waters et al., 2006; Chu et al., 2010).

Glutathione (GSH) is a multifunctional metabolite that has drawn extensive attention due to its unique structural properties, abundance, broad redox potential, and wide distribution in most living organisms. Along with ascorbate, GSH is considered as one of the most abundant redox couples in plant cells (Foyer and Halliwell, 1976). In plants, GSH is known to play a pivotal role in regulating stress responses as well as growth and development. Phenotypic analysis of GSH deficient Arabidopsis mutants demonstrated that GSH was essentially required for plant development, particularly embryo and meristem development (Vernoux et al., 2000; Cairns et al., 2006; Reichheld et al., 2007; Frottin et al., 2009; Bashandy et al., 2010). The *rml1* mutant, which was severely deficient in GSH, developed non-functional root meristem while the shoot meristem remained largely unaffected (Vernoux et al., 2000). In addition to serving as a source of reduced sulfur during secondary metabolite biosynthesis, GSH was widely reported to play crucial role in plant defense signalling network. Ball et al. (2004) reported that several stress responsive genes were altered due to changed GSH metabolism in *A. thaliana rax1-1* and *cad2-1* mutants of GSH biosynthesis enzyme. Another GSH deficient mutant, *pad2-1*, was demonstrated to be susceptible to *Pseudomonas syringae* as well as *P. brassicae* infections (Glazebrook and Ausubel, 1994; Glazebrook et al., 1997; Parisy et al., 2007; Datta and Chattopadhyay, 2015). Besides, GSH was shown to be involved in the modulation of NPR1-dependent and independent salicylic acid signalling pathways (Ghanta et al., 2011; Han et al., 2013). Subsequently it was reported that GSH induces ethylene biosynthetic pathway via transcriptional as well as post transcriptional regulation of the key enzymes (Datta et al., 2015).

Again, GSH exhibits a wide range of metal chelating activities and plays important role to reduce metal toxicity in plants. Detoxification of heavy metals within plant cell occurs via phytochelatins (PCs) that are synthesized from GSH by the enzyme phytochelatin synthase (Grill et al., 1989; Clemens et al., 1999; Vatamaniuk et al., 1999). On cadmium or copper exposure *A. thaliana* plants responded by increasing the transcription of *glutathione synthetase* and *glutathione reductase* (*GR*) genes which were involved in GSH synthesis and reduction respectively (Queval et al., 2009). Further, the involvement of GSH was reported in combating metal toxicity against arsenic stress in maize (Requejo and Tena, 2012), chromium toxicity in rice (Zeng et al., 2012; Qiu et al., 2013) and cadmium stress in *Pinus* and *A. thaliana* (Schützendübel et al., 2001; Jobe et al., 2012). Cross-talk of GSH with zinc and Fe homeostasis was also reported (Shanmugam et al., 2012). In another study, GSH was reported to trigger the upregulation of genes related to Fe uptake and transport and to increase the Fe concentration in *A. thaliana* seedlings under Fe deficiency (Koen et al., 2012). GSH-ascorbate redox cycle was studied against Fe deficiency as well (Ramírez et al., 2013). Previous studies demonstrated the association of GSH in maturation of Fe-S molecule and transport of dinitrosyl-Fe complexes in plants (Hider and Kong, 2011; Kumar et al., 2011). GSH also served as the reservoir of nitric oxide (NO) by formation of S-nitrosoglutathione (GSNO) complex which helped in NO mediated signalling including Fe deficiency responses (Chen et al., 2010; Ramirez et al., 2011). Subsequently, Shanmugam et al. (2015) revealed the role of GSH in enhancement of Fe deficiency tolerance in plants. The activities of several Fe responsive genes like *AtIRT1*, *AtFRO2* and *AtFIT* were reduced in *zir1*, a GSH depleted mutant of Arabidopsis, compared to the wild-type (WT). This observation attributed the role of GSH in regulation of Fe transport under Fe-limiting condition. However, the involvement of GSH in modulating subcellular Fe homeostasis in shoot has not been elucidated so far.

In this study, we report that under Fe deficient condition, GSH regulates subcellular Fe homeostasis by transcriptional induction of the vacuolar Fe exporters, *AtNRAMP3*, *AtNRAMP4* and the chloroplast Fe importer, *AtPIC1*. This GSH-mediated gene induction occurs via *S*-nitosylation of the Fe responsive basic helix loop helix (bHLH) transcription factors (TFs), bHLH29, bHLH38, and bHLH101.

## Materials and methods

### Plant growth, stress treatment and morphological analysis

The seeds of *A. thaliana* Columbia ecotype (Col-0) along with 2 GSH depleted mutants, *cad2-1* (N68137) and *pad2-1* (N3804), were procured from Nottingham Arabidopsis Stock Centre, UK. Seeds for the *noa1* (CS819466) and *gsnor1-3* (CS66012) mutants were procured from Arabidopsis Biological Resource Centre, USA. The *nramp3nramp4* double mutant that was generated by crossing the T-DNA insertion lines SALK_023049 for *nramp3-2* and SALK_085986 for *nramp4-3* as described by Molins et al. (2013) was used in this study. The seeds were inoculated in MS medium (Murashige and Skoog, 1962) after surface sterilization with 4 % sodium hypochlorite and Tween 20. The plants were maintained in a plant growth chamber at 21 °C with 60 % relative humidity and a photoperiod of 16 h light/8 h dark cycles (Datta et al., 2015).

The 7 d old seedlings were transferred to either minimal iron (MI) or depleted iron (DI) medium and maintained for 7 d. The MI medium contained 10 μM Fe in the MS medium. For preparation of DI medium, the Fe salt was omitted from MS medium keeping all other components unaltered. In addition, 300 µM ferrozine [3-(2-pyridyl)-5,6-diphenyl-1,2,4-triazine sulfonate] was also added to remove any trace amount of Fe from the medium (Eroglu et al., 2016). For control set, seedlings were maintained in MS medium for the entire period. After 7 d of stress treatment, the morphological parameters including primary root length, lateral root density and rosette diameter were measured using Image J software.

### RNA extraction and quantitative-RT PCR analysis

Total RNA was isolated from the tissue samples by Trizol method. Complementary DNA (cDNA) was prepared subsequently using iScript™ cDNA Synthesis Kit (Bio-rad) following manufacturer’s protocol. Quantitative PCR amplification was carried out in CFX96 Touch™ Real-Time PCR Detection System (Bio-rad) using iTaq™ Universal SYBR^®^ Green Supermix (Bio-rad) and gene specific primers (Supplementary Table S1). *AtActin2* was used as a reference gene to normalize the relative expression and *AtIRT1* was used as Fe-responsive marker gene to confirm the Fe deficient conditions.

### Chemical treatment of seedlings

14 d old seedlings were used for chemical treatments. For GSH feeding, freshly prepared 100 μM GSH was used while for BSO treatment, 1 mM BSO was used for 72 h as standardized before (Datta et al., 2015). For DTT treatment, a 5 mM DTT solution was used for 24 h treatment. In case of GSNO feeding, the seedlings were treated with 250 μ (Kailasham et al., 2018). For sodium nitroprusside (SNP) and cPTIO treatments, seedlings were treated with 1 µM SNP or 0.5 mM cPTIO respectively for 48 h (Ye et al., 2013). Control seedlings were maintained in half strength MS medium.

### Estimation of Fe content

Fe content from different tissues was measured through dry ash digestion method with slight modifications (Jiang et al., 2007). Briefly, tissues were harvested, washed with HPLC grade water, pat dried, weighed and used for ash preparation. The ash was digested with 0.5 M HNO_3_ and filtered through Whatman no. 42 filters. The solution was again filtered through Millex^®^ GV 0.22 μm PVDF membrane filter (Merck Millipore). The Fe content was analysed using ICP-OES (iCAP 6300 Duo ICP-OES, Thermo Scientific).

### Estimation of Chlorophyll content

The chlorophyll content was estimated following Lichtenthaler (1987). Briefly, 200 mg tissue was homogenized in 80 % acetone, followed by centrifugation at 5000 g for 5 min. The supernatant was used for the estimation of total chlorophyll.

### Estimation of total GSH content and GSH:GSSG ratio

GSH estimation was performed following Anderson (1985). Briefly, 200 mg of tissue was homogenized in 5 % sulphosalicylic acid followed by centrifugation at 12000 g for 20 min. GSH was estimated from the supernatant by 5,5’-dithiobis-(2-nitrobenzoic acid) (DTNB) method. For estimation of total GSH, 0.4 mM NADPH and 1 U GR were added to the reaction buffer and incubated for 20 min. Optical density was measured at 412 nm using a Double Beam UV-Vis Spectrophotometer (U2900, Hitachi). Amount of GSSG was calculated by subtracting GSH amount from total GSH.

### Isolation of chloroplast and vacuole

Chloroplast was isolated from leaves as standardized before (Kleffmann et al., 2004; Ghanta et al., 2014). Briefly, leaves were finely chopped in chloroplast extraction buffer (0.3 M sorbitol, 1 mM MgCl_2_, 50 mM HEPES/KOH, 2 mM EDTA, 0.04 % β-mercaptoethanol, 0.1 % PVPP, pH 7.8), filtered and centrifuged at 4 °C for 10 min at 1300 g. The pellet was suspended in isolation buffer (0.3 M sorbitol, 1 mM MgCl_2_, 50 mM HEPES/KOH, 2 mM EDTA, pH 7.8). The solution was loaded on the top of percoll gradient and centrifuged at 4 °C and 8000 g for 20 min. Intact chloroplasts were isolated and washed twice with the isolation buffer.

Vacuoles were isolated from leaves following Zouhar (2016). Briefly, leaves were collected, weighed, cut with razor blade into 2 mm strips and immersed in the protoplast isolation buffer [(1 % (w/v) Cellulase R10, 1 % (w/v) Macerozyme R10, 0.4 M mannitol, 25 mM β-mercaptoethanol, 10 mM 2-[N-morpholino]ethanesulfonic acid (MES)-KOH, (pH 5.7)]. The solution was then infiltrated and incubated for 4 h at room temperature with continuous shaking. The released protoplasts were filtered followed by centrifugation at 80 g at 20 °C for 15 min. 10 % Ficoll buffer was added to the pellet for protoplast disruption. The lysed protoplast solution was then treated with 4 % Ficoll buffer followed by vacuole isolation buffer (0.45 M mannitol, 5 mM sodium phosphate, 2 mM EDTA, pH 7.5). The solution was then ultracentrifuged at 50,000 g for 50 min at 10 °C. Vacuoles were collected from the 4 % Ficoll buffer/vacuole buffer interface.

The isolated chloroplast and vacuole fractions were used for Fe estimation as described above.

### Fe localization by Perls prussian blue and DAB staining

The Fe localization using Perls prussian blue staining technique was carried out following Rochhartdz et al. (2009). Briefly, the plant samples were fixed in a 6:3:1 methanol:chloroform:acetic acid (v/v) solution and washed twice with ultrapure water. The plants were then transferred to the pre-warmed staining solution (4 % (w/v) K_4_Fe(CN)_6_ and 4 % (v/v) HCl) and incubated at room temperature for 1 h. After removing the staining solution, the samples were washed twice with ultrapure water. For DAB intensification, the samples were incubated in preparation solution (0.01 M Na_2_CO_3_ and 0.3 % H_2_O_2_ in methanol) for 1 h. After washing with the 0.1 M phosphate buffer (pH 7) the samples were incubated in intensification solution (0.025 % (v/v) H_2_O_2_ and 0.005 % (w/v) CaCl_2_) and kept at room temperature for 30 min. The samples were then washed with ultrapure water and photographed.

### Vector construction and raising of transformed Arabidopsis plants

Genomic DNA was extracted from shoots following CTAB method. To clone the promoter regions of *AtPIC1*, *AtNRAMP3* and *AtNRAMP4* genes, approximately 1.5 kb of intergenic region upstream of the transcription start site was amplified by PCR using gene specific primers (Supplementary Table S1). The amplified promoter regions were cloned into pCAMBIA1304 vector between *BamHI* and *BglII* restriction enzyme (RE) sites to generate *AtNRAMP3pro*::*GUS and AtNRAMP4pro*::*GUS* constructs and between *BamHI* and *SpeI* to produce *AtPIC1pro*::*GUS* construct. These constructs were then transformed into Col-0 plants through *Agrobacterium* mediated floral dip transformation method (Clough and Bent, 1998). A vector control line harbouring the *CaMV35S::GUS* construct was generated as well. The transformed lines harbouring the recombinant constructs were selected and maintained up to T_2_ generations in a plant growth chamber as described above.

### Histochemical GUS assay

The seedlings from the transgenic lines harbouring the *AtPIC1pro*::*GUS, AtNRAMP3pro*::*GUS, AtNRAMP4pro*::*GUS* and *CaMV35S::GUS* constructs were incubated under DI condition or fed with GSH or GSNO as described above. The samples were then infiltrated with GUS staining solution (0.5 mg mL^−1^ 5-bromo-4-chloro-3-indolyl-β-D-glucuronic acid, 0.5 mM potassium ferrocyanide, 0.5 mM potassium ferricyanide, 0.1 % (v/v) Triton X-100, 100 mM phosphate buffer, pH 7.0, and 10 mM EDTA) following Jefferson et al. (1987). The stained samples were washed with 70 % ethanol and photographed.

### Yeast one-hybrid (Y1H) analysis

The 1456 bp upstream sequence from +1 site of *AtNRAMP3,* 1500 bp upstream sequence from +1 site of both *AtNRAMP4 and AtPIC1* genes were amplified by PCR and subsequently cloned into the *KpnI* and *SacI* RE sites of the *pAbAi* vector (Takara). On the other hand, the coding sequences (cds) of *bHLH29, bHLH38, bHLH39, bHLH100* and *bHLH101* gene were PCR amplified and cloned into pGADT7-Rec vector (Takara) following manufacturer’s protocol. The recombinant constructs was transformed into Y1H gold strain and Y1H assay was performed using Matchmaker^®^ Gold Yeast One-hybrid screening system (Takara) following the manufacturer’s protocol. The transformed yeast strains were grown on SD/-Leu and SD/–Leu/Aureobasidin-A media. The interaction between p53-pAbAi was considered as postive control while pAbAi alone was used as negative control.

### In silico promoter analysis and prediction of S-nitrosylation sites

The presence of bHLH TF-binding motifs was identified in the promoter regions of *AtNRAMP3*, *AtNRAMP4*, and *AtPIC1* using PlantPan 3.0 (plantpan.itps.ncku.edu.tw/promoter.php) and MEME suite 5.3.3 (https://meme-suite.org/; Bailey et al., 2009). The bHLH TFs like *At*bHLH29 (FIT), *At*bHLH38, and *At*bHLH101 were considered in this study and their protein sequences were retrieved from NCBI database. These sequences were submitted to GPS-SNO 1.0 software to predict the presence of *S*-nitrosylated cysteine residues (Xue et al., 2010).

### Protein stability assay

The coding sequences of *AtbHLH29, AtbHLH38* and *AtbHLH101* genes without stop codon was inserted into the vector *pVYCE* between *XbaI* and *SpeI* RE sites to obtain *35S::bHLH29-HA*, *35S::bHLH38-HA* and *35S::bHLH101-HA* constructs respectively. The recombinant constructs were transformed into *Agrobacterium tumifaciens* strain GV3101. Tobacco plants were infiltrated with the *Agrobacterium* strain harbouring the recombinant constructs and treated with 250 mM GSNO for 72 h. Next, the plants were treated with 200 µM cycloheximide and total proteins were isolated at 0, 3 and 6 h from GSNO treated and non Cl pH 7.5, 150 mM NaCl, 0.5 % Triton_X_100 (v/v) and protease inhibitor, Thermo Scientific) at 4 °C and centrifuged at 13,000 rpm for 30 min at 4°C. The supernatant was quantified by Bradford method and immunoblot analysis was performed using anti-HA antibody (1:1000 dilution, Sigma). Immunoblot using anti-αTubulin antibody (1:1000 dilution, Agrisera) was used as a control.

### In vivo 2,3-diaminonapthalene (DAN) assay of S-nitrosylated proteins

The fluorometric measurement of *S*-nitrosylated *At*bHLH29, *At*bHLH38 and *At*bHLH101 proteins were performed by DAN assay following Zhan et al. (2018) with slight modifications. Tobacco plants infiltrated with *Agrobacterium* strain harbouring the recombinant constructs *35S::bHLH29-HA*, *35S::bHLH38-HA* and *35S::bHLH101-HA* constructs were used in this experiment. 2 g of leaves from each of the infiltrated tobacco plants were homogenized in 3 ml lysis buffer (50 mM Tris, pH7.5, 150 mM NaCl, 10 mM MgCl_2_, 1 % glycerol, 0.5 % Triton X-100, 1mM PMSF as protease inhibitor). The sample was then centrifuged twice at 13000 g for 10 min at 4°C and the supermnatent was passed through Millex^®^ GV 0.22 μm PVDF membrane filter (Merck Millipore). Next, 80 µl anti-HA agarose beads (Thermo Scientific) were mixed with the lysates and incubated at 4^°^C for 2 h. The sample was then incubated with GSNO solution in a final concentration of 400 µM at dark for 30 min followed by rinsing with PBS buffer. The sample was then incubated with 300 µl of 200 µM of DAN and 200 µM of HgCl_2_ at dark for 30 min. The reaction was stopped with 2.8 M NaOH. The fluorescent signal of NAT compound of the sample was recorded in the Varioskan™ LUX multimode microplate reader (Thermo Scientific) using an excitation wavelength of 365 nm and emission wavelength of 450 nm. Next, the solution was used for determination of protein concentration using Bradford method. The DAN-NAT conversion rate of the sample was determined from the relative rluorescence intensity (RFU) of NAT molecule of per micromolar of protein.

### Statistical analysis

Statistical analysis of the data was performed using GraphPad Prism version 8.3.0 software (GraphPad Software, San Diego, California USA). The variation of morphological, and biochemical parameters as well as the relative transcript abundance among different samples were analyzed following one-way ANOVA or two-way ANOVA followed by Dunnett’s multiple comparison tests. The statistical significance at P ≤ 0.05 was considered to determine the difference between two sets of data. Details of statistical tests, replicates, sample size, and the significance levels of P-values were indicated in respective figure legends. The data were represented as mean ± standard error of mean (SEM).

## Results

### GSH played a crucial role in regulating Fe transporters under Fe deficiency

Since GSH is known to play an important role in regulating Fe deficiency signaling, the response of the GSH depleted mutants, *cad2-1* and *pad2-1*, to Fe deficient conditions were analyzed. The MS medium containing 100 μM Fe was used as a control medium while the MI medium containing 10 μM Fe and the DI medium lacking any Fe source and containing the Fe-chelator, ferrozine were used as Fe deficient media. Since, the expression of *AtIRT1* gene was found to be the highest after 7 d treatment, this time point was used for all downstream analyses (Figure S1). The mutant lines when grown under MI and DI conditions displayed increased sensitivity to Fe deficiency as compared with the Col-0 plants (Figure 1, S2, S3).

**FIGURE 1.**
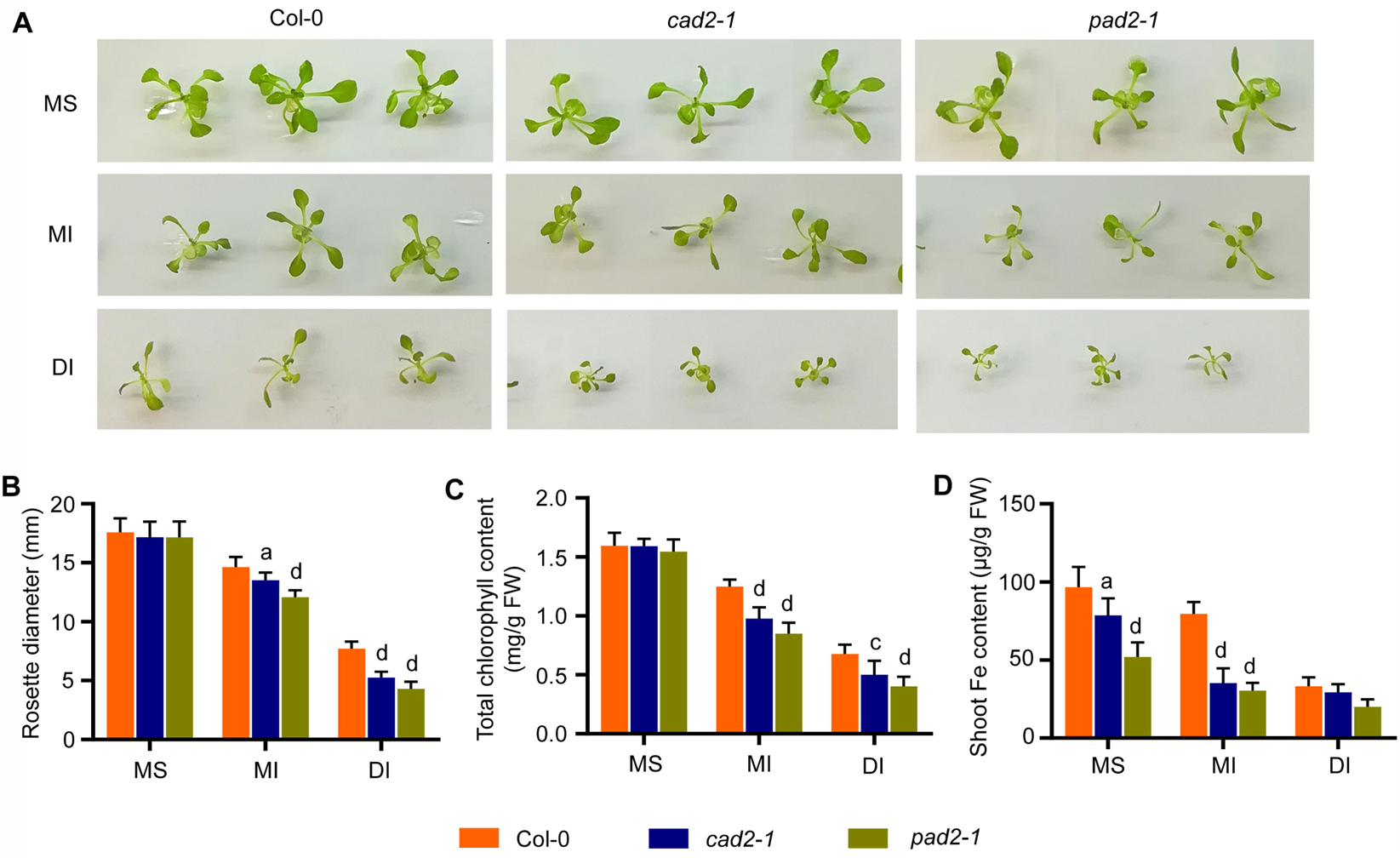
Response of GSH depleted mutants under Fe deficient conditions. 7 d old MS grown seedlings of Col-0, *cad2-1* and *pad2-1* were exposed to MI and DI conditions for 7 d and analyzed. (A) Shoot morphology, (B) rosette diameter, (C) total chlorophyll content, and (D) shoot Fe content. Results were represented as mean±SEM (n=9). Statistical differences between the genotypes were analyzed by two-way ANOVA followed by Dunnett’s multiple comparison test and denoted by different alphabets at P<0.05 (a), P< 0.001 (c), and P< 0.0001 (d).

Since the GSH depleted mutants were sensitive to Fe deficiency, the next pertinent point was to identify the Fe transporters that might be regulated by GSH. To this end, the expression of the Fe transporters and several genes involved in regulating Fe homeostasis were analyzed in the mutant lines and compared with the Col-0 plants. Interestingly, it was found that the expression of *AtNRAMP3*, *AtNRAMP4*, *AtIRT1*, *AtYSL8*, *AtFER1*, *AtFRO3*, and *AtFRO7* genes were significantly down-regulated in the *pad2-1* mutant while *AtPIC1* was significantly down-regulated in both the mutants (Figure 2A, B). On the other hand, the expression of *AtYSL1* and *AtYSL3* genes were found to be up-regulated in the *pad2-1* mutant. Among these differentially expressed genes, *AtNRAMP3* and *AtNRAMP4* are known to be involved in vacuolar Fe efflux while *AtPIC1* encodes a chloroplastic Fe importer. This observation suggested a probable involvement of GSH in modulating subcellular Fe homeostasis in plants. We, therefore, checked the expression of these three genes under DI condition. It was found that under DI condition, the relative transcript abundance for all these genes were significantly lower in both the mutant lines further supporting our previous observations (Figure 2C).

**FIGURE 2.**
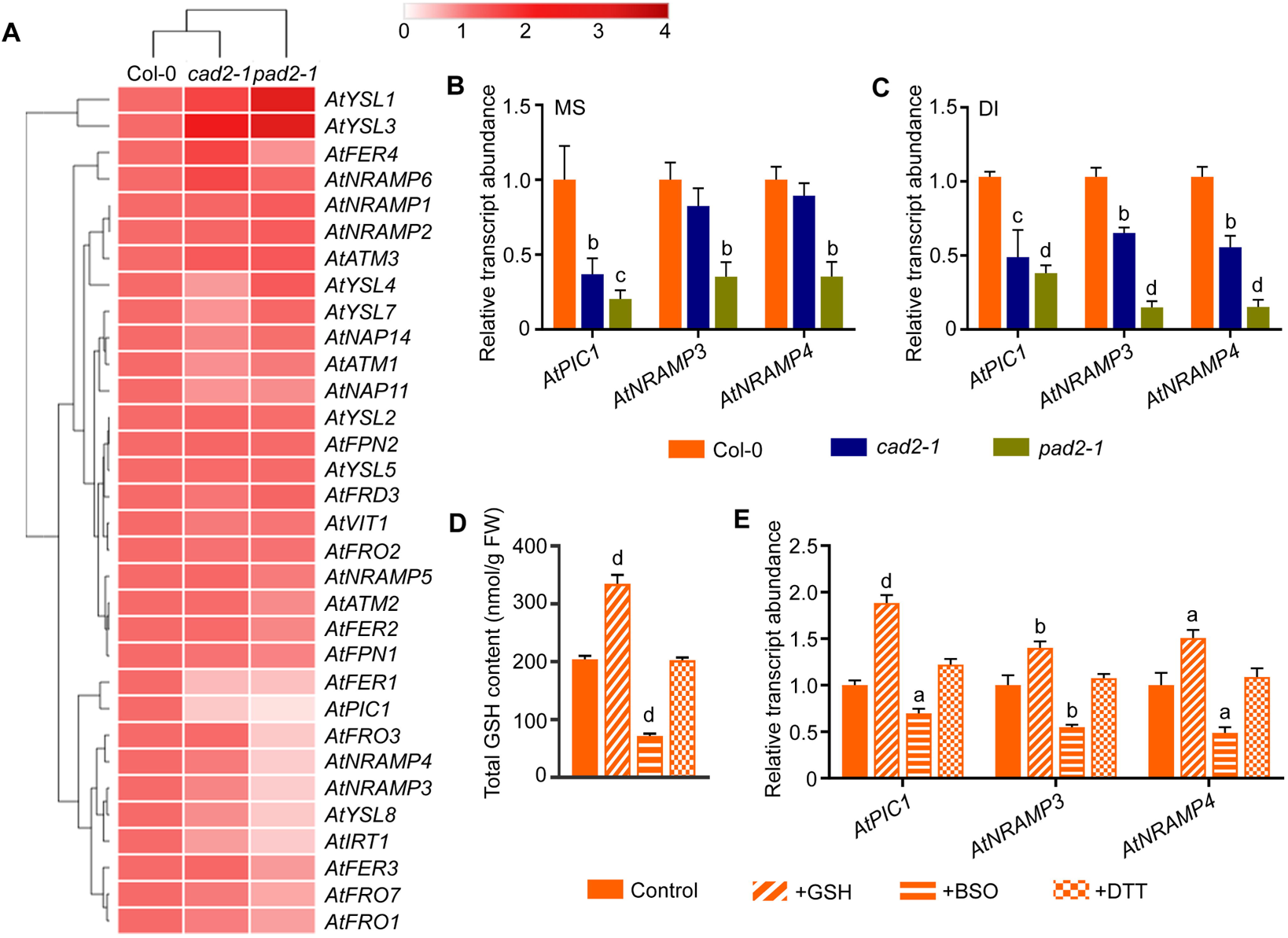
Expression analyses of different Fe transporters under altered GSH conditions. (A) 14 d old MS grown Col-0, *cad2-1*, and *pad2-1* plants were analyzed for the expression pattern of various Fe responsive genes and results were depicted as heat map. Relative transcript abundance of *AtPIC1*, *AtNRAMP3*, and *AtNRAMP4* genes were analyzed from Col-0, *cad2-1*, and *pad2-1* plants grown under (B) MS and (C) DI conditions. 14 d old MS grown Col-0 plants were treated with 100 μM GSH, or 1 mM BSO, or 5 mM DTT solutions and analyzed. Control plants were maintained in half strength MS medium for the entire duration. (D) Total GSH content and (E) relative transcript abundance of *AtPIC1*, *AtNRAMP3*, and *AtNRAMP4* genes. Results were represented as mean±SEM (n=3). Statistical differences between the genotypes or treatments were analyzed by one-way ANOVA followed by Dunnett’s multiple comparison test and denoted by different alphabets at P<0.05 (a), P<0.01 (b), P< 0.001 (c), and P< 0.0001 (d).

### Exogenously altered GSH level was sufficient to regulate expression of the Fe transporters

To further confirm the GSH-mediated regulation of the identified Fe transporter genes, Col-0 seedlings were exogenously treated with GSH or an inhibitor of GSH biosynthesis, BSO. In addition, a non-specific reducing agent DTT was also used for treatment. A half strength MS medium was used as a control set. To confirm the efficiency of the feeding treatments, total GSH content was estimated from each set. The total GSH level was found to be significantly increased in the GSH-fed plants, decreased in the BSO-fed plants while no significant alteration was observed in the DTT-fed plants (Figure 2D). Next, the relative transcript abundance of the identified genes was analyzed. It was observed that the expression of the *AtNRAMP3*, *AtNRAMP4*, and *AtPIC1* genes were induced in response to GSH feeding while the expression was decreased in response to BSO treatment (Figure 2E). The DTT feeding, however, failed to significantly alter the expression levels thus indicating that non-specific reducing condition was not sufficient for this regulation.

Since the *pad2-1* mutant is severely deficient in GSH content, we supplemented these mutant seedlings with exogenous GSH or DTT. It was observed that only GSH supplementation, and not DTT feeding, could compensate for its deficiency and resulted in an increased expression of the identified Fe transporters (Figure S4).

### GSH modulated subcellular Fe homeostasis under Fe depleted condition

The Fe transporters *AtNRAMP3* and *AtNRAMP4* are vacuolar exporters for Fe while the *AtPIC1* is responsible for Fe influx into the chloroplast. Since GSH was found to regulate the expression of all of these genes, it was hypothesized that GSH might be involved in modulating the subcellular Fe homeostasis under Fe deficient condition. To dissect this, the organellar Fe content was estimated from the *cad2-1* and *pad2-1* mutants along with the Col-0 plants under both control and DI conditions. It was observed that the chloroplast Fe content was lower in the mutant lines as compared with the Col-0 plants under control as well as DI conditions (Figure 3A). On the other hand, the vacuolar Fe content was significantly lower in the mutants under control condition but higher under DI condition as compared with the Col-0 plants (Figure 3B). This observation suggested that the mutant lines with depleted GSH levels failed to efficiently export the Fe from vacuoles and channel them into the chloroplast under DI condition. To further validate this hypothesis, the chloroplast Fe content was estimated from the *nramp3nramp4* double mutant under DI condition. It was observed that exogenous GSH treatment under DI condition could improve the chloroplast Fe content in Col-0 plants while it failed to do so in the *nramp3nramp4* double mutant (Figure 3C).

**FIGURE 3.**
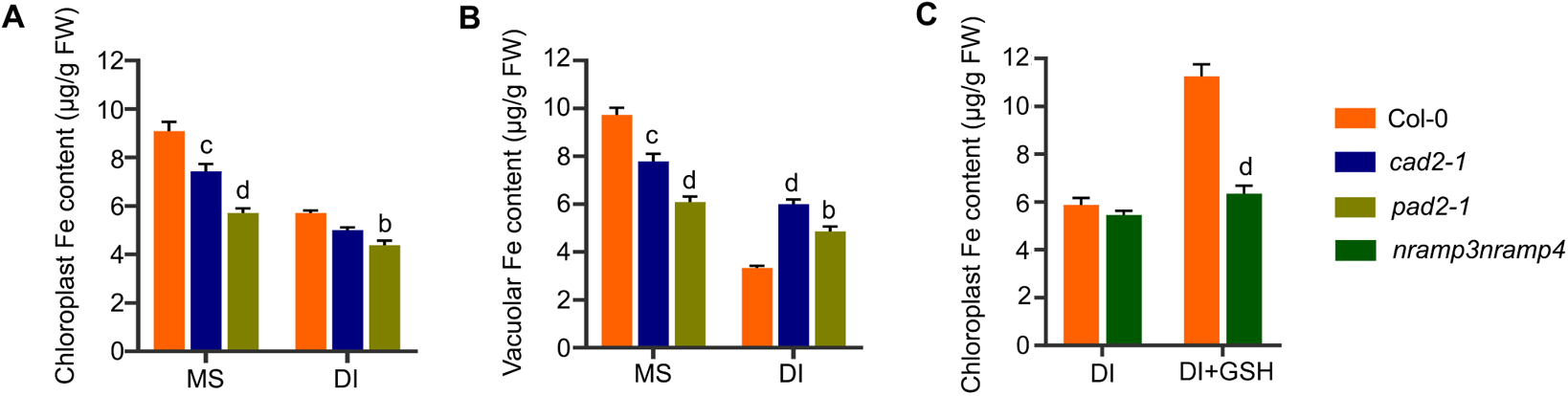
Estimation of organellar Fe contents under MS and DI conditions. 7 d old MS grown Col-0, *cad2-1* and *pad2-1* seedlings were exposed to DI condition for 7 d. Control plants were maintained in the MS medium for the entire period. Chloroplasts and vacuoles were isolated from the samples and used for Fe content estimation. (A) Chloroplast Fe content, and (B) vacuolar Fe content. Results were represented as mean±SEM (n=3). Statistical differences between the genotypes were analyzed by two-way ANOVA followed by Dunnett’s multiple comparison test and denoted by different alphabets at P<0.01 (b), P< 0.001 (c), and P< 0.0001 (d). (C) Col-0 and *nramp3nramp4* double mutant was similarly exposed to DI condition with or without 100 μM GSH and the chloroplast Fe content was analyzed. Results were represented as mean±SEM (n=3). Statistical differences between the genotypes were analyzed by two-way ANOVA followed by Sidak’s multiple comparison tests and denoted by alphabet at P< 0.0001 (d).

### GSH-mediated regulation of Fe homeostasis involved GSNO

Since the association of GSH with GSNO is known in regulating Fe deficiency response, we were curious if this GSH-mediated modulation of subcellular Fe homeostasis also involved GSNO. Therefore, the Col-0 and *pad2-1* plants were treated with GSH, GSNO and SNP in combination with their inhibitors and DI condition. The expression of *AtNRAMP3*, *AtNRAMP4*, and *AtPIC1* genes were then analyzed. When Col-0 plants were treated with GSH, GSNO or the NO donor, SNP under DI condition, stronger induction of the genes was observed as compared with DI condition alone (Figure 4A, B, C). Treating the plants with SNP along with the GSH inhibitor, BSO under DI condition did not display this effect. Yet again, plants treated with GSH in combination with the NO inhibitor, cPTIO failed to induce the gene expression even under DI condition. These observations suggested that the GSH mediated regulation of the identified Fe transporter genes occurred in a GSNO-dependent manner. Similarly, in case of *pad2-1* mutant, the treatment with GSH or GSH-cPTIO combination under DI condition showed a similar trend like that of Col-0 plants. On contrary, GSNO or SNP treatment under DI condition failed to augment gene expression (Figure 4D, E, F).

**FIGURE 4.**
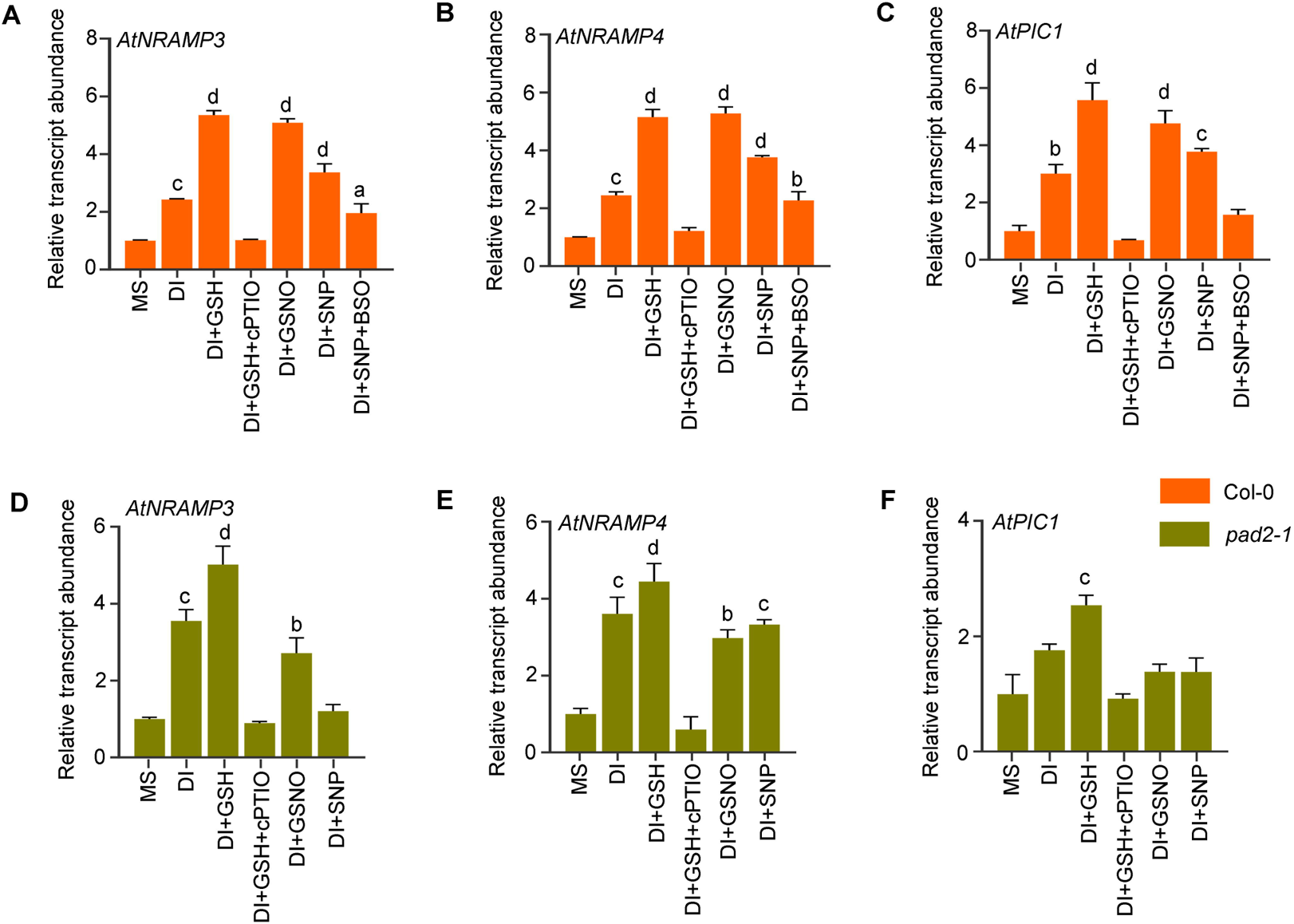
Expression analyses of the identified Fe transporter genes in response to chemical treatments. 7 d old Col-0 and *pad2-1* seedlings were transferred to DI condition for 7 d with or without GSH or GSNO or SNP or GSH/cPTIO or SNP/BSO combinations. For control set, plants were maintained in half strength MS medium for the entire duration. Shoot samples collected from each set were used for qRT-PCR analysis of (A, D) *AtNRAMP3*, (B, E) *AtNRAMP4*, and (C, F) *AtPIC1* genes. Results were represented as mean±SEM (n=3). Statistical differences between the treatments were analyzed by one-way ANOVA followed by Dunnett’s multiple comparison test and denoted by different alphabets at P<0.05 (a), P<0.01 (b), P< 0.001 (c), and P< 0.0001 (d).

To gain a deeper insight, the expression of the identified Fe transporters were analyzed in the NO deficient mutant, *noa1* and the *gsnor1-3* mutant that accumulates higher level of GSNO (Figure 5). Interestingly, it was observed that the expression of all three genes was significantly down-regulated in the *noa1* mutant as compared to the Col-0 plants under DI condition. On the other hand, the expression of the *AtNRAMP4* and *AtPIC1* genes were found to be significantly higher in the *gsnor1-3* mutant while *AtNRAMP3* displayed similar transcript abundance as that of the Col-0 plants. This observation strongly suggested the involvement of GSNO in regulating these Fe transporter genes.

**FIGURE 5.**
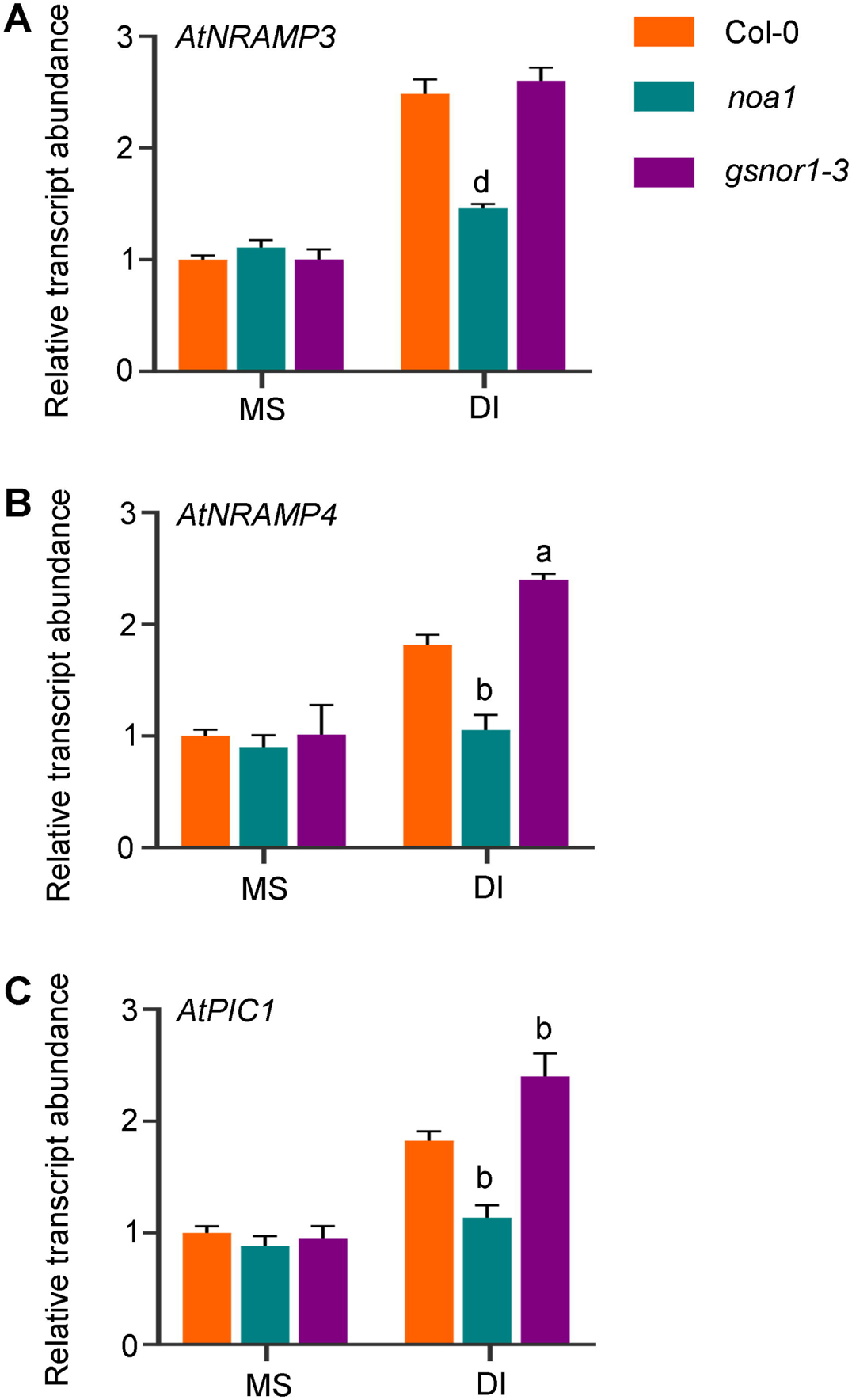
Expression analyses of the identified Fe transporter genes in the *noa1* and *gsnor1-3* mutants. 7 d old Col-0, *noa1* and *gsnor1-3* seedlings were exposed to DI condition for 7 d. For control set, plants were maintained in MS medium for the entire duration. Shoot samples collected from each set were used for qRT-PCR analysis of (A) *AtNRAMP3*, (B) *AtNRAMP4*, and (C) *AtPIC1* genes. Results were represented as mean±SEM (n=3). Statistical differences between genotypes were analyzed by two-way ANOVA followed by Dunnett’s multiple comparison test and denoted by different alphabets at P<0.05 (a), P<0.01 (b), and P< 0.0001 (d).

### GSH triggered promoter activation of the Fe transporter genes

Since GSH modulated the relative transcript abundance for the identified Fe transporter genes, it was assumed that a probable transcriptional regulation might be involved. To identify the mechanism of this regulation, transgenic lines harboring the *AtNRAMP3pro::GUS*, *AtNRAMP4pro::GUS,* and *AtPIC1pro::GUS* constructs were generated (Figure S5). A transgenic line harboring the *CaMV35S::GUS* construct was used as a negative control. The promoter activities in response to GSH and GSNO treatments were analyzed by histochemical GUS assay. The DI condition was used as a positive control for these treatments. The GUS activity in all the transgenic lines was found to be induced in response to both GSH and GSNO treatments which was also supported by the relative transcript abundance of the *GUS* gene (Figure 6). On contrary, the GUS activity as well as its expression was unaltered in the *CaMV35S::GUS* containing negative control line. This observation strongly suggested that the GSH-mediated regulation of these Fe transporters occurred via transcriptional activation.

**FIGURE 6.**
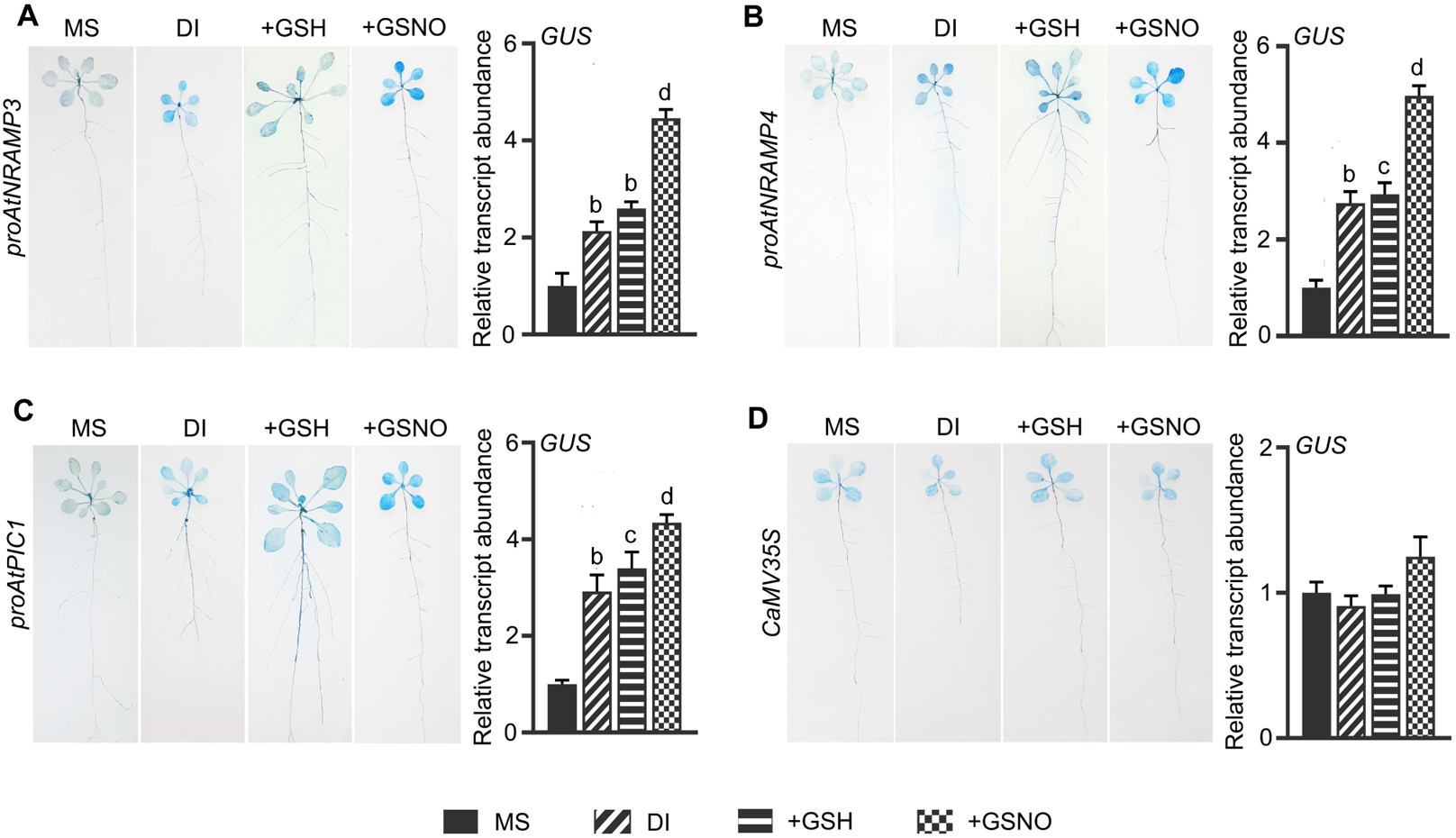
Promoter analyses of the identified Fe transporter genes. The seedlings of different transgenic lines were exposed to GSH or GSNO treatment and used to analyze their promoter activity by histochemical GUS assay and qRT-PCR analysis for *GUS* gene expression. The DI condition was used as a positive control. (A) *proAtNRAMP3* (harbouring *AtNRAMP3pro::GUS* construct), (B) *proAtNRAMP4* (harbouring *AtNRAMP4pro::GUS* construct), and (C) *proAtPIC1* (harbouring *AtPIC1pro::GUS* construct). A control transgenic line harbouring the *CaMV35S::GUS* construct was used as a negative control (D). Representative images of histochemical GUS staining were presented. The experiment was independently repeated thrice. For relative transcript abundance, results were represented as mean±SEM (n=3). Statistical differences between the treatments were analyzed by one-way ANOVA followed by Dunnett’s multiple comparison test and denoted by different alphabets at P<0.01 (b), P< 0.001 (c), and P< 0.0001 (d).

### GSH mediated transcriptional induction involved S-nitrosylation of Fe-responsive TFs

Several bHLH TFs including *At*FIT (AtbHLH29), *At*bHLH38, *At*bHLH39, *At*bHLH100, and *At*bHLH101 were known to be involved in the transcriptional regulation of Fe responsive genes, *AtIRT1* and *AtFRO2* in roots (Gao et al., 2019; 2020). This prompted us to explore if the *AtNRAMP3*, *AtNRAMP4* and *AtPIC1* genes were also regulated by these bHLH TFs. The bHLH TFs were reported to bind at the conserved E-box DNA binding (CANNTG) motif (Toledo-Ortiz et al., 2003). Therefore, the promoter regions of *AtNRAMP3*, *AtNRAMP4*, and *AtPIC1* genes were analyzed for the presence of this conserved CANNTG motif. Interestingly, the promoters of all these identified Fe transporter genes contained the conserved E-box motif indicating their probable interaction with the bHLH TFs (Figure 7A, B, Table S2). With this clue, Y1H analysis was performed to check if these bHLH TFs can regulate the transcription of the *AtNRAMP3*, *AtNRAMP4*, and *AtPIC1* genes. It was identified that *At*bHLH38 can bind to the *AtNRAMP3* promoter, *At*bHLH38 and *At*bHLH101 to the *AtNRAMP4* promoter while *At*bHLH29 and *At*bHLH101 can interact with the *AtPIC1* promoter (Figure 7C). Next, to check if these TFs were regulated in a GSNO-dependent fashion their amino acid sequences were screened for probable *S*-nitrosylation sites using the GPS-SNO 1.0 software. Among them, *At*bHLH29, *At*bHLH38 and *At*bHLH101 were found to contain 3, 4, and 2 putative *S*-nitrosylation sites respectively (Figure 7D; Table S3). To further confirm these *in silico* data, the effect of GSNO treatment on the stability of these TFs was analyzed. Interestingly, it was observed that the stability of these three bHLH TFs was enhanced by GSNO treatment (Figure 7E). Moreover, the DAN assay was performed to analyze the DAN-NAT conversion by the NO released from the *S*-nitosylated thiol group present in these proteins and compared with that of GSNO which has a single *S*-nitrosylated Cys residue. It was observed that the DAN-NAT conversion ratio was ∼2 fold higher in case of *At*bHLH29 while similar in case of *At*HLH38 and *At*bHLH101 as that of GSNO (Figure 7F). This result indicated that *At*bHLH29 contains at least 2 *S*-nitrosylated Cys residues while *At*bHLH38 and *At*bHLH101 posses atleast 1 *S*-nitrosylated Cys residue each. Together, these observations suggested that the GSH-mediated regulation of the identified Fe transporters involved transcriptional modulation via these *S*-nitrosylated bHLH TFs.

**FIGURE 7.**
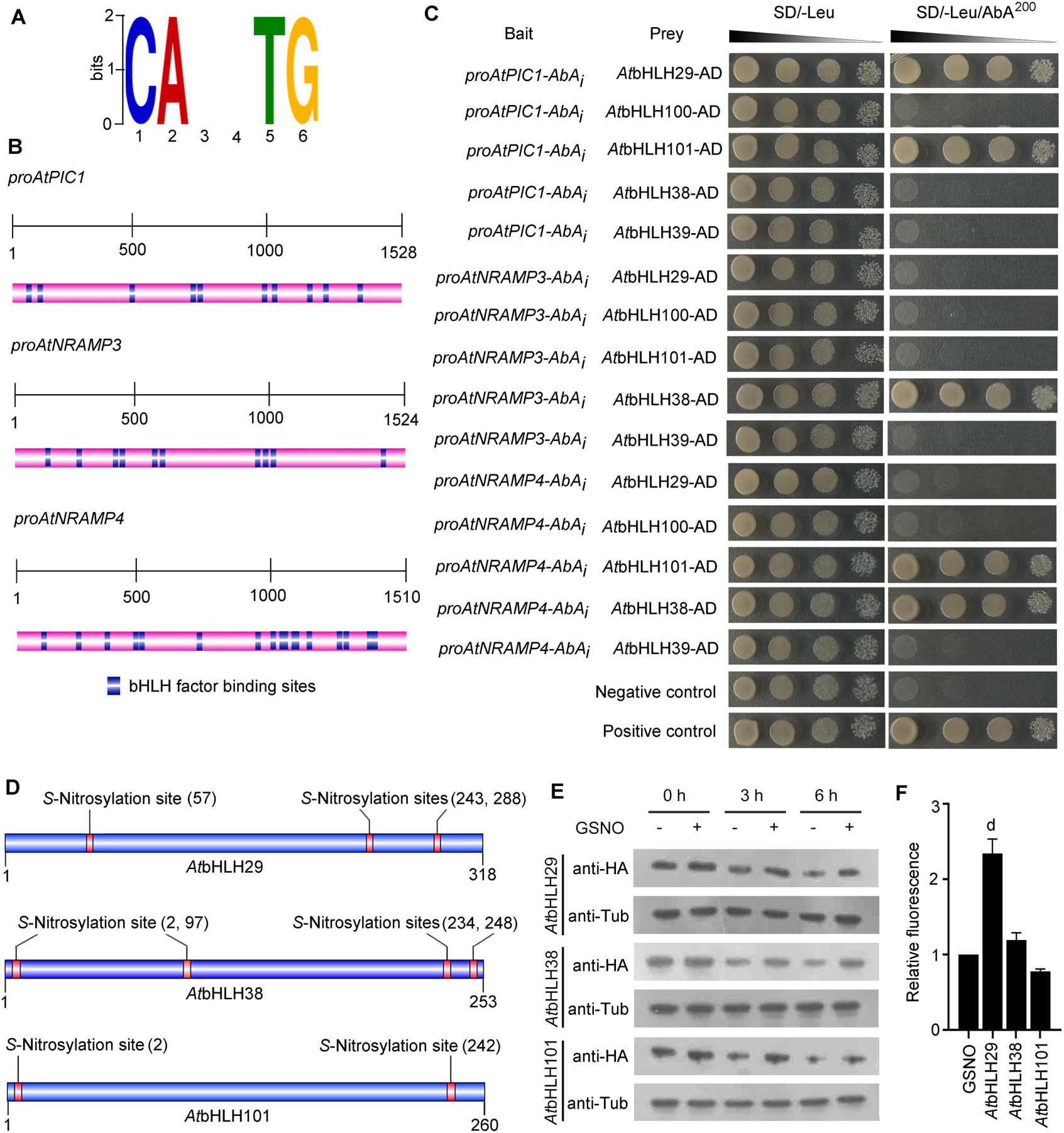
*S*-Nitrosylation of Fe responsive bHLH TFs regulate transcription of the identified Fe transporters. *In silico* promoter analysis was performed for the identified Fe transporters using PlantPAN 3.0 and MEME suite 5.3.3. (A) bHLH factor binding motif, and (B) presence of bHLH factor binding motifs in the promoter sequences of the *AtNRAMP3*, *AtNRAMP4*, and *AtPIC1* genes. (C) Y1H analysis for interaction of the Fe responsive bHLH TFs, *At*bHLH29, *At*bHLH38, *At*bHLH39, *At*bHLH100, and *At*bHLH101 with the *AtNRAMP3*, *AtNRAMP4*, and *AtPIC1* promoters. (D) *In silico* analysis for identification of putative *S*-nitrosylation sites in the *At*bHLH29, *At*bHLH38, and *At*bHLH101 TFs using GPS-SNO 1.0 software. (E) Protein stability assay in response to exogenous GSNO treatment for the *At*bHLH29, *At*bHLH38, and *At*bHLH101 TFs using anti-HA antibody. Immunoblot using anti-αTubulin (anti-αTub) antibody was used as a control. (F) DAN assay of *S*-nitrosylated Cys residues in the *At*bHLH29, *At*bHLH38, and *At*bHLH101 TFs. The protein stability assay and the DAN analysis were independently repeated thrice. For, DAN assay, results were represented as mean±SEM (n=3). Statistical differences between the sets were analyzed by one-way ANOVA followed by Dunnett’s multiple comparison tests and denoted by alphabet at P< 0.0001 (d).

## Discussion

In the last few decades the role of GSH in regulating multiple stress responses in plants was elaborately studied. Recently, emerging evidences suggested a positive role of GSH in mitigating Fe deficiency responses in plants (Koen et al., 2012; Ramirez et al., 2013). It was demonstrated that during Fe deficiency in *Arabidopsis*, the elevated level of GSH could significantly regulate different Fe responsive genes like *FRO*2, *IRT1*, *NAS4*, and *FIT1* to enhance Fe uptake and transport through roots (Koen et al., 2012). Subsequently, the GSH mediated regulation of these genes was found to occur in a GSNO dependent manner (Garcia et al., 2010, Shanmugam et al., 2015; Kailasam et al., 2018). However, the role of GSH in regulating subcellular Fe homeostasis in plants was not elucidated so far. In our previous study, comparative proteomic analysis of *pad2-1*, a GSH depleted mutant, revealed the down-accumulation of *At*FER1 protein in shoot as compared with the Col-0 plants (Datta et al., 2015). This clue prompted us to investigate the role of GSH in regulating the expression of different Fe responsive genes in *Arabidopsis* shoot. To begin with, we selected two GSH depleted mutants *cad2-1* and *pad2-1*, and analyzed their response under Fe limited conditions. Both the mutants displayed severe sensitivity to Fe deficiency as compared with the Col-0 plants (Figure 1). This observation was in line with the earlier study where Fe deficiency responses were found to be aggravated by reduced GSH content in *Arabidopsis* (Ramirez et al., 2013; Shanmugam et al. 2015). The next approach was to identify the Fe transporters and homeostasis related genes that might be responsible for this observation. Out of the 32 Fe transporters analyzed, 2 vacuole Fe exporters, *AtNRAMP3* and *AtNRAMP4*, and a chloroplast Fe importer, *AtPIC1* were found to be significantly down-regulated under GSH deficient condition, while 2 chloroplast Fe exporters, *AtYSL1* and *AtYSL3* were up-regulated (Figure 2).

The altered expression of these organellar Fe transporters suggested a possible role of GSH in modulating the subcellular Fe homeostasis under Fe limited condition. The vacuole, during Fe sufficient condition stores the excess amount of Fe to rescue the cell from Fe mediated oxidative damage. This stored Fe is released under Fe limited condition to mitigate Fe deficiency responses in plants (Morrissey and Guerinot, 2009). On contrary, *AtPIC1*, a chloroplast Fe importer, was found to be up-regulated during Fe deficiency to maintain the Fe homeostasis in chloroplast (Duy et al., 2007). In a previous study, it was reported that *AtNRAMP3* and *AtNRAMP4* function as major vacuolar Fe exporters in germinating seeds where vacuolar reserve is the primary source of Fe (Bastow et al., 2018). Further, it was demonstrated that the Fe mobilization into chloroplast was limited when the vacuolar Fe could not be retrieved. In this study, GSH was found to positively regulate the expression of *AtNRAMP3*, *AtNRAMP4*, and *AtPIC1* genes. This made us inquisitive if GSH can help in channeling the Fe retrieved from vacuoles into the chloroplast in shoot under Fe deficient conditions. We, therefore, analyzed the Fe content in the vacuoles and the chloroplasts in the GSH depleted mutants. Surprisingly, both the vacuolar and chloroplast Fe contents were found to be lower in the mutants under Fe sufficient condition (Figure 3). This lower Fe accumulation in the mutants can be attributed to their impaired Fe uptake due to reduced *AtIRT1* expression in the roots. On contrary, the vacuolar Fe content was found to be higher in the mutants while the chloroplast Fe content remained lower under DI condition. This observation strongly suggested that the GSH depleted mutants failed to efficiently retrieve the vacuolar Fe under DI condition and channel them into the chloroplast. Further supporting this observation, under DI condition, exogenous GSH treatment could improve chloroplast Fe content in Col-0 plants while it failed to do so in the loss-of-function *nramp3nramp4* double mutant.

GSH combines with NO leading to the formation of GSNO which acts as an important signaling intermediate. Earlier, GSNO was reported to act as a key modulator of Fe responsive genes in Arabidopsis (Koen et al., 2012; Shanmugam et al. 2015; Kailasam et al., 2018). This made us interested to dissect the role of GSNO in regulating the identified Fe transporter genes. It was observed that under DI condition, exogenous feeding with GSH, GSNO, or SNP resulted in a stronger induction of these genes indicating a positive role of GSH-GSNO module in regulating these genes. Again, the NO scavenger cPTIO could nullify the effect of GSH feeding while the GSH biosynthesis inhibitor BSO reversed the effect of SNP treatment further supporting this hypothesis (Figure 4). Under DI condition, the expression of these transporter genes was again significantly down-regulated in the *noa1* mutant further supporting this observation (Figure 5). Together, these observations strongly suggested that GSH regulated the subcellular Fe homeostasis via a GSNO mediated pathway.

The next pertinent question was how this GSH-GSNO module induced the expression of the *AtNRAMP3*, *AtNRAMP4*, and *AtPIC1* genes. Interestingly, promoter analysis through histochemical GUS assay confirmed that all the 3 genes were induced in response to GSH or GSNO treatments via transcriptional activation (Figure 6). Since neither GSH nor GSNO can directly bind to these promoters, this transcriptional activation presumably involved one or more TFs. Earlier reports have identified several bHLH TFs that are involved in regulating various Fe responsive genes in *Arabidopsis* (Yuan et al., 2008; Gao et al., 2019; 2020). Among them, *At*bHLH29 (FIT1, clade IIIa) and clade Ib bHLH factors (*At*bHLH38, *At*bHLH39, *At*bHLH100, *At*bHLH101) were reported to regulate two crucial Fe responsive genes, *AtIRT1* and *AtFRO2* while these TFs were, in turn, regulated by yet other bHLH TFs (Colangelo and Guerinot, 2004; Yuan et al., 2008; Gao et al., 2019). However, the TFs involved in regulating *AtNRAMP3*, *AtNRAMP4*, and *AtPIC1* genes were not reported till date. We observed that the promoters of these genes contain conserved E-box motifs indicating their probable transcriptional regulation via bHLH TFs. Further, the *At*bHLH29, *At*bHLH38, and *At*bHLH101 TFs were identified to regulate the transcription of these transporter genes (Figure 7).

Next, we wanted to understand how these bHLH TFs were modulated by the GSH-GSNO module. The GSNO mediated regulation is often known to involve *S*-nitrosylation of the TFs (Darbani et al., 2013). We were curious if these bHLH TFs could be regulated by GSNO as well and analyzed for the presence of putative *S*-nitrosylation sites. Interestingly, multiple *S*-nitrosylation sites were detected in all these three TFs and their protein stability was found to be significantly enhanced in response to GSNO treatment (Figure 7). These observations strongly suggested a GSH mediated regulation of these bHLH TFs via *S*-nitrosylation which corroborated with the earlier report where *At*FIT1 stability was reduced in presence of the NO scavenger cPTIO (Meiser et al., 2011).

In summary, it can be postulated that during Fe deficiency the accumulation of GSH in cells can activate the vacuolar Fe exporters like *AtNRAMP3* and *AtNRAMP4* to facilitate Fe export from the vacuolar reserve. On the other hand, GSH was found to trigger the expression of the chloroplast Fe importer, *AtPIC1* to maintain chloroplast Fe content during Fe deficiency in plants (Figure 8). This regulation involved GSH-GSNO mediated transcriptional activation of these transporter genes via *S*-nitrosylation of the Fe responsive TFs, *At*bHLH29, *At*bHLH38 and *At*bHLH101.

**FIGURE 8.**
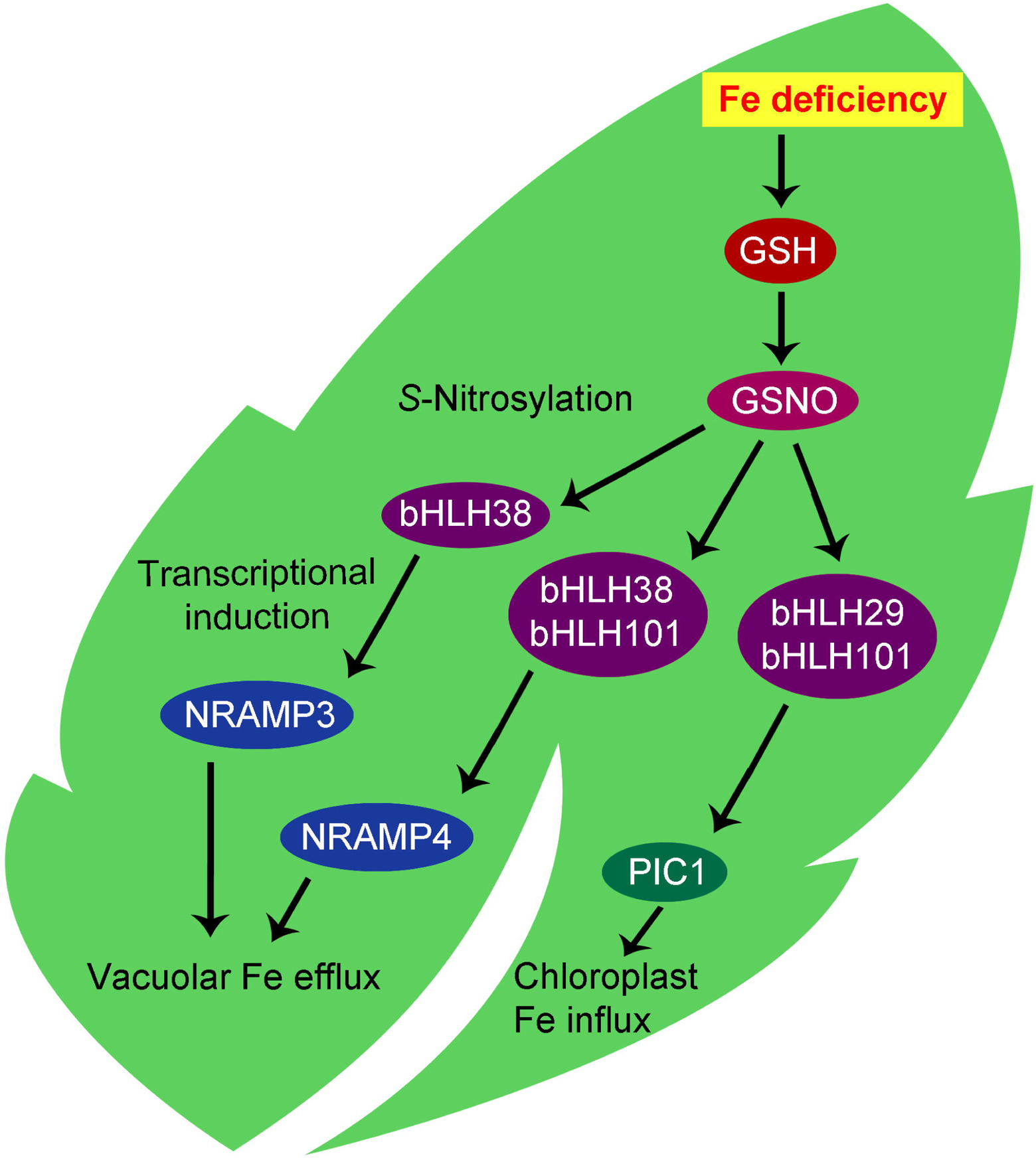
Model for GSH mediated regulation of subcellular Fe homeostasis under Fe limited condition. Fe deficiency leads to the accumulation of GSH in cells. GSH then activates the vacuolar Fe exporters, *At*NRAMP3 and *At*NRAMP4, to facilitate release of Fe from the vacuolar reserve. GSH also induces the chloroplast Fe importer, *At*PIC1 which aids in channelling Fe into the chloroplast. This GSH mediated modulation involves transcriptional induction via GSNO by *S*-nitrosylation of the Fe responsive bHLH TFs, *At*bHLH29, *At*bHLH38, and *At*bHLH101.

## Author Contributions

RD and SP conceived and designed the research plan; RS raised the transgenic lines, performed expression analysis, feeding experiments and GUS assay; SG standardized the plant growth and stress treatments, PK performed biochemical experiments, SS generated the constructs and performed Y1H analysis; CR helped in maintaining the transgenic lines and performed protein stability and DAN assays, DS performed *in silico* analysis; RD and SP analyzed the data and prepared the manuscript.

## Acknowledgement

We thank the central instrumentation facility of the Department of Botany, University of Calcutta and Department of Botany, Dr A. P. J. Abdul Kalam Government College. We thank Prof. Manoj Prasad, NIPGR, New Delhi, India for providing us the pCAMBIA1304 vector. We are thankful to Prof. Sebastien Thomine, Institut de Biologie Integrative de la Cellule, CNRS, France and Prof. Janneke Balk, John Innes Centre, Norwich, UK for sharing the *nramp3nramp4* double mutant with us.

## Supplementary data

Figure S1. Expression of *AtIRT1* gene in Col-0 shoot under DI condition.

Figure S2. Biochemical analyses of Col-0, *cad2-1* and *pad2-1* plants in response to Fe deficient conditions.

Figure S3. Perls-DAB staining of Col-0, *cad2-1* and *pad2-1* plants grown under MS condition.

Figure S4. Response of *pad2-1* plants to exogenous GSH and DTT treatments.

Figure S5. Screening of transgenic lines harbouring (A) *CaMV35S::GUS*, (B) *AtPIC1pro::GUS*, (C) *AtNRAMP3pro::GUS*, and (D) *AtNRAMP4pro::GUS* constructs.

Table S1. List of primers used

Table S2. *In sillico* analysis of *AtNRAMP3*, *AtNRAMP4* and *AtPIC1* promoters for the presence of bHLH TF binding motifs

Table S3. Prediction of putative *S*-nitrosylation sites using GPS-SNO 1.0 software

