## Supplementary Figure for "Glutathione regulates transcriptional activation of iron transporters via *S*-nitrosylation of bHLH factors to modulate subcellular iron homeostasis"

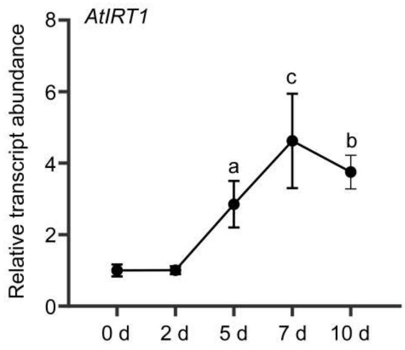


**Figure S1. Expression of *AtIRT1* gene in Col-0 shoot under DI condition.** 7 d old MS grown seedlings were transferred to DI medium and samples were collected after 0 d, 2 d, 5 d, 7 d, and 10 d of treatment. Results were represented as mean±SEM (n=3). Statistical differences between different time points were denoted by different alphabets at P<0.05 (a), P<0.01 (b), and P< 0.001 (c).

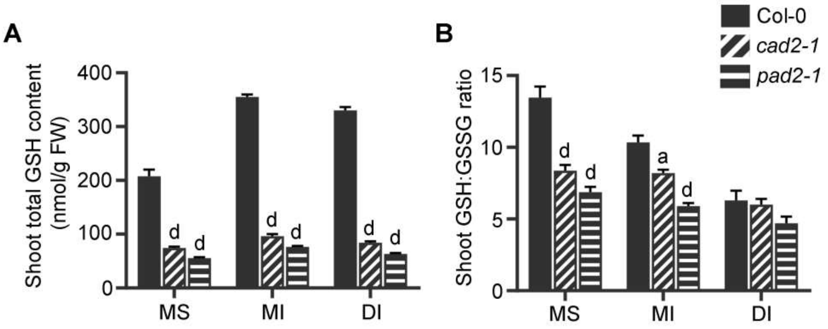


**Figure S2. Biochemical analyses of Col-0, *cad2-1* and *pad2-1* plants in response to Fe deficient conditions**. 7 d old MS grown seedlings were exposed to MI and DI condition for 7 d and biochemical parameters were recorded. (A) Total GSH content, and (B) GSH:GSSG ratio. Results were represented as mean±SEM (n=3). Statistical differences between the Col-0, *cad2-1* and *pad2-1* were denoted by different alphabets at P<0.05 (a) and P< 0.0001 (d).

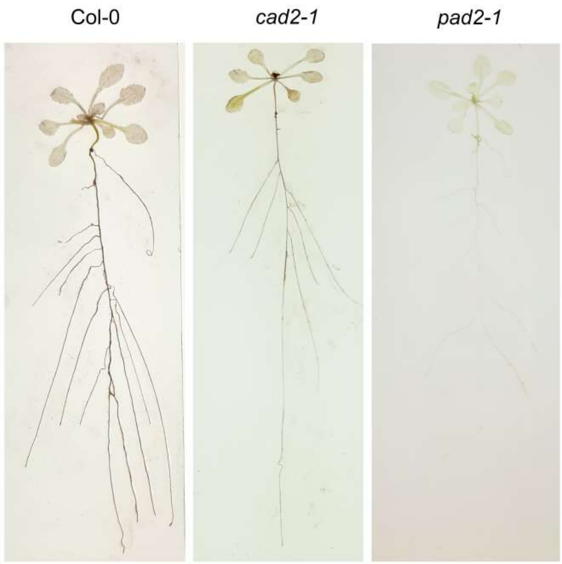


**Figure S3. Perls-DAB staining of Col-0, *cad2-1* and *pad2-1* plants grown under MS condition.**

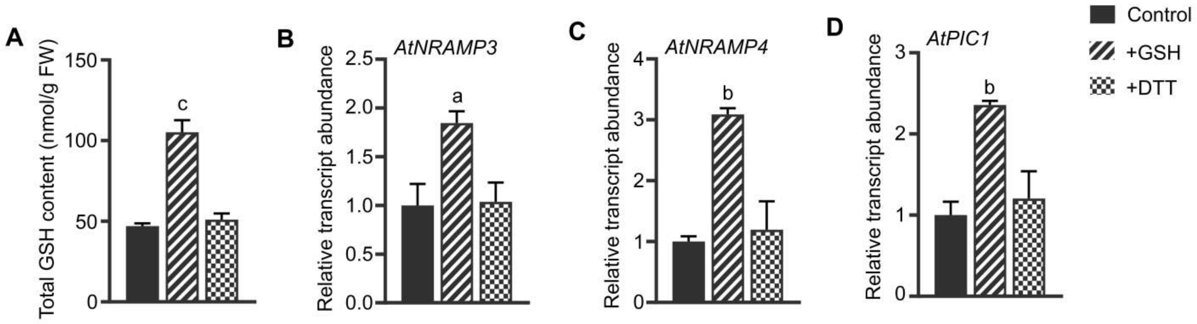


**Figure S4. Response of *pad2-1* plants to exogenous GSH and DTT treatments.** (A) Total GSH content. (B-D) Relative transcript abundance of the identified Fe transporter genes. Results were represented as mean±SEM (n=3). Statistical differences between the treatments were denoted by different alphabets at $P<0.05$ (a), $P<0.01$ (b), and $P< 0.001$ (c).

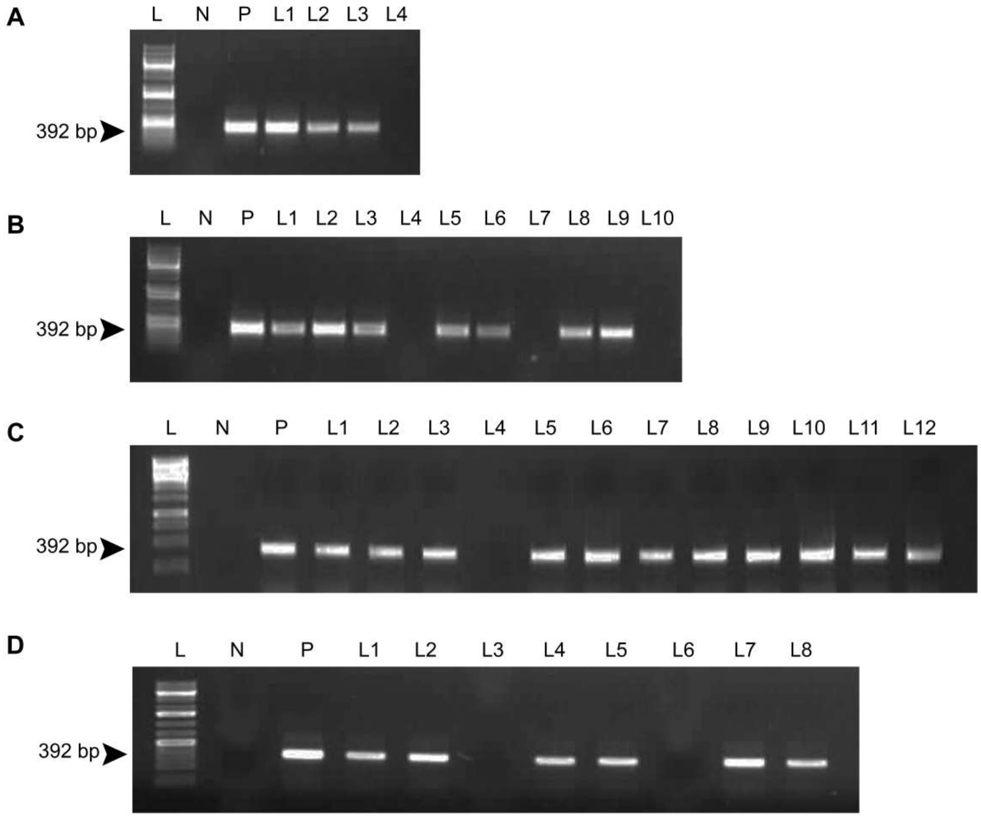


**Figure S5. Screening of transgenic lines harbouring (A) *CaMV35S::GUS*, (B) *AtPIC1pro::GUS*, (C) *AtNRAMP3pro::GUS*, and (D) *AtNRAMP4pro::GUS* constructs.**
