## Supplementary Table for "Glutathione regulates transcriptional activation of iron transporters via *S*-nitrosylation of bHLH factors to modulate subcellular iron homeostasis"

Table S1: List of primers used

| Accession No | Gene name | Primer sequence (5’-3’) |
|---|---|---|
| NM_100041 | *AtFRO1* | F: TACCCACCTCTGGTCTCACC |
| | | R: TGATCGATTGGGTGAATGTG |
| AY302057.1 | *AtFRO2* | F: AAAGCAATAACGGTGGTTCG |
| | | R: CCCAAACAAGCTACGACCAT |
| NM_001332575.1 | *AtFRO3* | F: GGACCGGTATCTCGCATTTA |
| | | R: CACAAGGAAGAACACGAGCA |
| NM_124352.4 | *AtFRO7* | F: TGCAGCTGAGTTTCTTGGAA |
| | | R: TGACCAAGCCAAACATGGTA |
| U27590.1 | *AtIRT1* | F: TGGTGTTGGAGCTCCTCTCT |
| | | R: TACCAACTGCGTTCTTGCTG |
| AF165125.1 | *AtNRAMP1* | F: ACGCTGATTCTTCTCGCACT |
| | | R: AAACCTGCAAGCTTCCTTGA |
| AF141204.1 | *AtNRAMP2* | F: CCGGCTTAAGAAATGGATGA |
| | | R: TACAAGTGCAGCCACAGTCC |
| AF202539.1 | *AtNRAMP3* | F: TGGAGTTGTGGGTTGCATTA |
| | | R: CACACCGCCTCCATATTTCT |
| AF202540.1 | *AtNRAMP4* | F: TTTGCTTGGATGTTTGGTCA |
| | | R: GAAAGAAACCGCAAGAGCAC |
| NM_117995.2 | *AtNRAMP5* | F: CGGCGAATTCAAACGTCTAT |
| | | R: CGCCCATAGAAGAAGCCATA |
| NM_101464.4 | *AtNRAMP6* | F: GGTTATGCCGCACAATCTCT |
| | | R: AGCTCCATTTACCCACAACG |
| NM_118544.4 | *AtYSL1* | F: TCTGCTGTTGCTTGTTACGG |
| | | R: TGTGTGTGGAAGCCATTGAT |
| NM_122346.3 | *AtYSL2* | F: GGAGTGAACACCGAAGGAAA |
| | | R: CCAAAGAATGCCCAAAAGAA |
| NM_001203603.2 | *AtYSL3* | F: TGGTTCCCATCAACTCTTCC |
| | | R: GCGATGATCGAGACAACAGA |
| NM_123465.4 | *AtYSL4* | F: GTACCCGAGTGGAAGGAACA |
| | | R: CGCTATAAGCAAGGCCGTAG |
| NM_112646.2 | *AtYSL5* | F: ATCTATATCTGCGCGCCTGT |
| | | R: CACACAACCCATTGCTGTT |
| NM_113616.3 | *AtYSL6* | F: TCGCTTCCGTTGTAGGAAGT |
| | | R: TCACTGCGTAAGGTGCTTTG |
| NM_105247.4 | *AtYSL7* | F: TCAAAGTCTTAGGCCGCACT |
| | | R: GCGTTGCAAAAGGCTAAGAC |
| NM_103733.4 | *AtYSL8* | F: TCGAGAAGCTCTCTTGCACA |
| | | R: TGATGCTAGCGACCAATCAG |
| AF229850.1 | *AtFER1* | F: GGTGGACACAAACAACATGC |
| | | R: TCTCAGCATGCCCTCTCTCT |
| NM_111942.4 | *AtFER2* | F: TTCTGAATCTGCCATCAACG |
| | | R: CAAAGACAATGCAAGCTCCA |
| NM_115467.5 | *AtFER3* | F: AAGGGTCTCGCCAAGTTTTT |
| | | R: TGGACATCATCGTTCTTGGA |
| NM_129588.5 | *AtFER4* | F: ATAACATCGCGCTCAAAGGT |
| | | R: TGCCAAGTGGACATCATTGT |

| | | |
|---|---|---|
| NM_121413.2 | *AtNAP14* | F: CGTCCTCCTGGAACACAGTT |
| | | R: CCTTGGCCAACCAAAAGTAA |
| NM_126238.3 | *AtVIT1* | F: ATCATTACGCGAGGGAGATG |
| | | R: CGGTATAAAACCGCCAAGAA |
| NM_119005.3 | *AtATM1* | F: ATGAGGGGATCTCGGTTTCT |
| | | R: CGTGCGGAGGATCTTCTTAG |
| NM_119004.1 | *AtATM2* | F: TGATGCTGAGTGGTGGAGAG |
| | | R: ATTTCCCCAACAACACTTCG |
| NM_125212.3 | *AtATM3* | F: CTGTTGATTGGCTTGCTTCA |
| | | R: CGCCTGTTTCTCGACTTAGG |
| NM_001336703.1 | *AtFPN1* | F: AACAACGCGAAGAAGGAGAA |
| | | R: CTGGAACAGGAGCCAGAGTC |
| NM_001342716.1 | *AtFPN2* | F: TTGTGCTCCCTGGAGTTTCT |
| | | R: CTCCGGTTCTGAGAGGTGAG |
| NM_127089.4 | *AtPIC1* | F: ATATCAATCGCCACCGTCTC |
| | | R: AATCACAGCTGCAACCACAG |
| NM_111683.2 | *AtFRD3* | F: CTCCAGCTCCGGATACAAAG |
| | | R: AATCCACGAAAGATGCCTTG |
| NM_105215.3 | *AtNAP11* | F: GTGTAAAGAGCGAGGGCTTG |
| | | R: CCCTTGTCTGGAGCAAGAAG |
| NM_001338359.1 | *AtActin2* | F: GCACCCTGTTCTTCTTACCG |
| | | R: AACCCTCGTAGATTGGCACA |
| AF202539.1 | *AtNRAMP3pro* | F: GGATCCATAAAACGGGACGGGTTGGC |
| | | R: AGATCTCGGGTTTTGTTTTCGAGG |
| AF202540.1 | *AtNRAMP4pro* | F: GGATCCCAATCAGATGGAGCCCTACA |
| | | R: AGATCTTTCCCGGTAACCGCCGATTC |
| NM_127089.4 | *AtPIC1pro* | F: GGATCCTCTTGCCGCCGCTTAGTGAA |
| | | R: ACTAGTGAGTAAAGTAGCGAATTGGTG |
| CP073591.1 | *GUS (uidA)* | F: CCCTTACGCTGAAGAGATGC |
| | | R: GGCACAGCACATCAAAGAGA |
| AF202539.1 | *AtNRAMP3* (for pABAi) | F: GAGCTCATAAAACGGGACGGGTTGGC |
| | | R: GGTACCCGGGTTTTGTTGTTTTCGAGG |
| AF202540.1 | *AtNRAMP4* (for pABAi) | F: GAGCTCCAATCAGATGGAGCCCTACA |
| | | R: GGTACCTTCCCGGTAACCGCCGATTC |
| NM_127089.4 | *AtPIC1* (for pABAi) | F: GAGCTCTCTTGCCGCCGCTTAGTGAA |
| | | R: GGTACCGAGTAAAGTAGCGAATTGGTG |
| NM_115556.4 | *AtbHLH38* (for pGADT7-Rec) | F: AATGTGTGCATTAGTCCCTTCA |
| | | R: GAATTAGTCACCTAGTTAAACGA |
| NM_115557.3 | *AtbHLH39* (for pGADT7-Rec) | F: ATGTGTGCATTAGTACCTCCAT |
| | | R: CTCGACTACAACGCTTTGTC |
| NM_129689.2 | *AtbHLH100* (for pGADT7-Rec) | F: ATGTGTGCACTTGTCCCTCC |
| | | R: TCATGTAAACGAGTGTCCACAT |
| NM_120497.2 | *AtbHLH101* (for pGADT7-Rec) | F: ATGTGTACCTTAACGCCAATGT |
| | | R: TTATGATTGGCGTAATCCCAAGAG |
| FJ410331.1 | *AtbHLH29* (for pGADT7-Rec) | F: ATGGAAGGAAGAGTCAACGC |
| | | R: CAATCTCGGTTACATCATCAC |
| NM_115556.4 | *AtbHLH38* (for pVYCE) | F: TCTAGAAATGTGTGCATTAGTCCCTTCA |
| | | R: ACTAGTGAATTAGTCACCTAGTTAAACGA |

| | | |
|---|---|---|
| NM_120497.2 | *AtbHLH101* (for pVYCE) | F: TCTAGAATGTGTACCTTAACGCCAATGT |
| | | R: ACTAGTTTATGATTGGCGTAATCCCAAGAG |
| FJ410331.1 | *AtbHLH29* (for pVYCE) | F: TCTAGAATGGAAGGAAGAGTCAACGC |
| | | R: ACTAGTCAATCTCGGTTACATCATCAC |

Table S2: *In sillico* analysis of *AtNRAMP3*, *AtNRAMP4* and *AtPIC1* promoters for the presence of bHLH TF binding motifs

| Matrix ID | Conserved motif | Promoter | Position | Strand | Similarity score | Hit sequence |
|---|---|---|---|---|---|---|
| TFmatrixID_0174 | 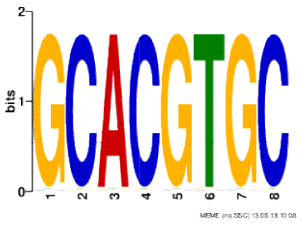 | *AtNRAMP3* | 466 | + | 0.75 | CCACGagc |
| | | | 466 | - | 0.75 | ccaCGAGC |
| | | *AtNRAMP4* | 1002 | + | 0.75 | GCGCGtgg |
| | | | 1002 | - | 0.75 | gcgCGTGG |
| | | | 1231 | + | 0.75 | ACACGttc |
| | | | 1231 | - | 0.75 | acaCGTTC |
| | | *AtPIC1* | 36 | - | 0.75 | gctCTTGC |
| | | | 97 | + | 0.75 | GCACCggc |
| TFmatrixID_0497 | 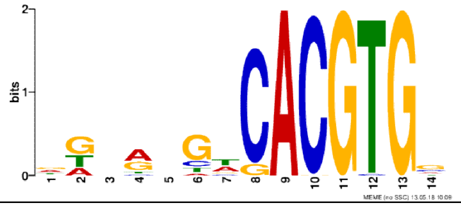 | *AtNRAMP3* | 450 | + | 0.89 | ggtgagagACGTGt |
| | | *AtNRAMP4* | 1100 | - | 0.88 | cCACGTcgcatttt |
| TF_motif_seq_0300 | 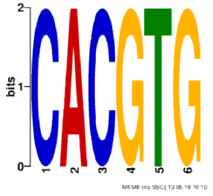 | *AtNRAMP3* | 457 | + | 0.83 | GACGTg |
| | | | 457 | - | 0.83 | gACGTG |
| | | | 467 | + | 0.83 | CACGAg |
| | | | 467 | - | 0.83 | cACGAG |
| | | | 595 | + | 0.83 | CACCTg |
| | | | 595 | - | 0.83 | cACCTG |
| | | | 971 | + | 0.83 | CACGAg |
| | | | 971 | - | 0.83 | cACGAG |
| | | | 993 | + | 0.83 | TACGTg |
| | | | 993 | - | 0.83 | tACGTG |
| | | | 1442 | + | 0.83 | AACGTg |
| | | | 1442 | - | 0.83 | aACGTG |
| | | *AtNRAMP4* | 1003 | + | 0.83 | CGCGTg |
| | | | 1003 | - | 0.83 | cGCGTG |

| | | | | | | |
|---|---|---|---|---|---|---|
| | | | 1046 | + | 0.83 | CACGTt |
| | | | 1046 | - | 0.83 | cACGTT |
| | | | 1101 | + | 0.83 | CACGTc |
| | | | 1101 | - | 0.83 | cACGTC |
| | | | 1232 | + | 0.83 | CACGTt |
| | | | 1232 | - | 0.83 | cACGTT |
| | | | 1307 | + | 0.83 | GACGTg |
| | | | 1307 | - | 0.83 | gACGTG |
| | | *AtPIC1* | 1246 | + | 0.83 | CAGGTg |
| | | | 1246 | - | 0.83 | cAGGTG |
| TF_motif_seq_0301 | 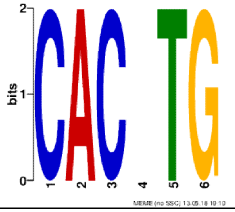 | *AtNRAMP3* | 595 | + | 1 | CACCTg |
| | | *AtPIC1* | 1246 | - | 1 | cAGGTG |
| TF_motif_seq_0302 | 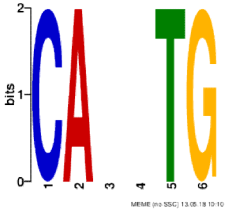 | *AtNRAMP3* | 595 | + | 1 | CACCTg |
| | | | 595 | - | 1 | cACCTG |
| | | *AtNRAMP4* | 110 | + | 1 | CAGATg |
| | | | 110 | - | 1 | cAGATG |
| | | | 275 | + | 1 | CATTTg |
| | | | 275 | - | 1 | cATTTG |
| | | | 500 | + | 1 | CATATg |
| | | | 500 | - | 1 | cATATG |
| | | | 916 | + | 1 | CAATTg |
| | | | 916 | - | 1 | cAATTG |
| | | | 1385 | + | 1 | CAGTTg |
| | | | 1385 | - | 1 | cAGTTG |
| | | *AtPIC1* | 497 | + | 1 | CATTTg |
| | | | 497 | - | 1 | cATTTG |
| | | | 1199 | + | 1 | CATTTg |
| | | | 1199 | - | 1 | cATTTG |
| | | | 1246 | + | 1 | CAGGTg |

| | | | | | | |
|---|---|---|---|---|---|---|
| | | | 1246 | - | 1 | cAGGTG |
| | | | 1430 | + | 1 | CAACTg |
| | | | 1430 | - | 1 | cAACTG |
| TF_motif_seq_0410 | 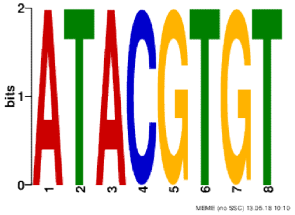 | *AtNRAMP3* | 119 | + | 0.75 | ATACAtat |
| | | | 119 | - | 0.75 | ataCATAT |
| | | | 238 | - | 0.75 | aaaGGTAT |
| | | | 456 | + | 0.88 | AGACGtgt |
| | | | 456 | - | 0.75 | agaCGTGT |
| | | | 458 | - | 0.75 | acgTGTAT |
| | | | 631 | + | 0.75 | ATACAtgc |
| | | | 929 | + | 0.75 | ATACAtat |
| | | | 929 | - | 0.75 | ataCATAT |
| | | | 992 | + | 0.88 | ATACGtga |
| | | *AtNRAMP4* | 399 | + | 0.75 | ATAAGttt |
| | | | 497 | - | 0.88 | acaCATAT |
| | | | 704 | - | 0.75 | gcaCATAT |
| | | | 1023 | + | 0.88 | ATTCGtgt |
| | | | 1045 | + | 0.75 | ACACGttt |
| | | | 1045 | - | 0.88 | acaCGTTT |
| | | | 1173 | + | 0.75 | ATACTttt |
| | | | 1281 | - | 0.75 | tcaCATAT |
| | | *AtPIC1* | 787 | + | 0.75 | ATAGGcgt |
| | | | 789 | - | 0.75 | aggCGTAT |
| | | | 1001 | - | 0.75 | aaaCCTAT |
| TFmatrixID_0112 | 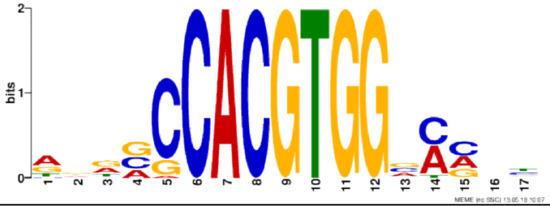 | *AtNRAMP4* | 1096 | + | 0.85 | ttcacCACGTcgcat tt |
| TFmatrixID_0159 | 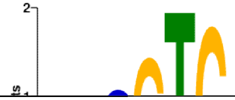 | *AtNRAMP4* | 1100 | - | 0.98 | cCACGTc |

| | | | | | | |
|---|---|---|---|---|---|---|
| | | | 1307 | + | 0.98 | gACGTGg |
| TFmatrixID_0169 | 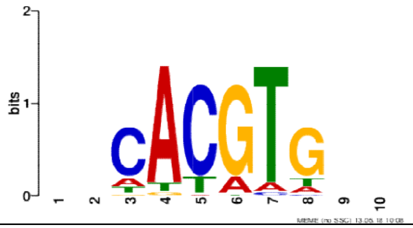 | *AtNRAMP4* | 1230 | + | 0.92 | taCACGTtct |
| TFmatrixID_0177 | 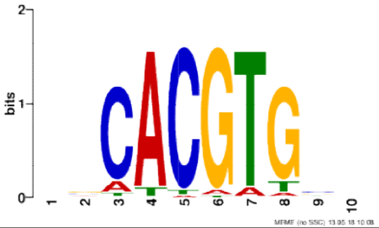 | *AtNRAMP4* | 1230 | + | 0.89 | taCACGTtct |
| TFmatrixID_0559 | 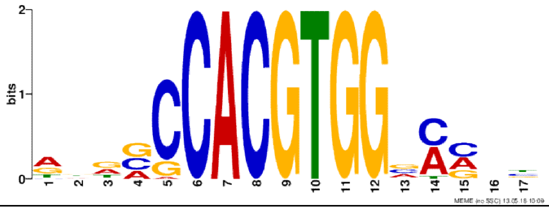 | *AtNRAMP4* | 1099 | + | 0.87 | acCACGTcgc |
| TFmatrixID_0815 | 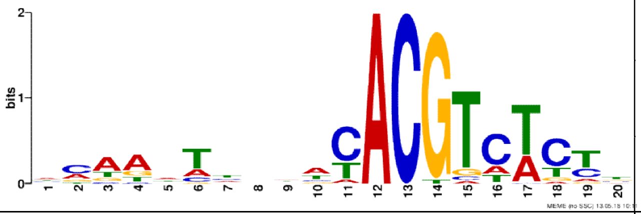 | *AtNRAMP4* | 1036 | + | 0.93 | ccaggccaaaCACGTttcta |
| | | | 1091 | + | 0.87 | ttctgttcacCACGTcgcat |
| | | | 1303 | - | 0.9 | ttctgACGTGgcgtaatagt |
| TFmatrixID_0823 | 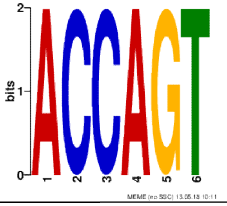 | *AtPIC1* | 1049 | - | 1 | aCTGGT |

S3: Prediction of putative S-nitrosylation sites using GPS-SNO 1.0 software

| TF | Position | Peptide | Score |
|---|---|---|---|
| *At*bHLH29 | 57 | GPLQNSPCFIDENQF | 19.577 |
| | 243 | GFYVRLVCNKGEGVA | 0.918 |
| | 288 | TYTLDGTCFEQSLNL | 2.201 |
| *At*bHLH38 | 2 | ******MCALVPSFF | 27.788 |
| | 97 | LFSSLRSCLPASDQS | 2.245 |
| | 234 | MDDYKINCEELSERM | 1.304 |
| | 248 | MLYLYEKCENSFN** | 1.59 |
| *At*bHLH101 | 2 | ******MCTLTPMFP | 28.036 |
| | 242 | HLQMRGDCKVRLEEL | 18.818 |
